# Precision Aptamers Against a Native GPCR through Ligand-Guided Selection

**DOI:** 10.64898/2026.05.04.722743

**Authors:** Agbor otu Egbe Vydaline, Mitali Bhate, Divyani Sitaldin, Yilam Ng Cen, Sergei Rozhkov, German Sosa, Amedee des Georges, Prabodhika Mallikaratchy

## Abstract

G protein–coupled receptors (GPCRs) constitute the largest and most diverse class of membrane receptors encoded in the human genome. They detect a wide range of chemical and physical stimuli and transduce these signals into intracellular responses through highly regulated pathways. Reflecting their central role in physiology, GPCRs are among the most prominent targets in drug discovery. However, identifying ligands that recognize GPCRs in their native conformational and membrane context remains a significant challenge. Here, we report an expanded aptamer discovery platform based on ligand-guided selection (LIGS) to isolate aptamers against GPCRs in their native cellular state. Using the β_2_-adrenergic receptor (β_2_AR) as a model system and employing agonists and antagonists as competing ligands, we identified three aptamers with high specificity for β_2_AR. These aptamers exhibit selective binding to cell-surface β_2_AR, showing higher apparent affinity towards cell-membrane bound β_2_AR than toward the purified receptor, which is consistent with recognition of native receptor context. Beyond target recognition, we show that the selected aptamers induce rapid internalization, indicating functional engagement. Together, these findings establish ligand-guided selection as a generalizable strategy for the discovery of conformationally sensitive aptamers targeting GPCRs in their native membrane environments.

**Significance:** The ability to discover ligands for receptors that undergo dynamic conformational changes is essential for advancing targeted therapeutics. G protein–coupled receptors (GPCRs), among the most sought-after drug targets, exist in transient and heterogeneous conformational states that are difficult to replicate in purified or artificial systems. Here, we introduce a ligand discovery platform that leverages native receptor interactions with agonists and antagonists to enable the selection of nucleic acid ligands (aptamers) directly against GPCRs in their cellular context, eliminating the need for purified receptors. The resulting aptamers exhibit selective binding to membrane-bound receptors and display intracellular functionality, highlighting a broadly applicable strategy for discovering ligands that recognize and modulate GPCRs in their native environments.

## Introduction

G protein–coupled receptors (GPCRs) represent the largest and most diverse class of membrane receptors with more than 800 members identified in the human genome^1,2^. GPCRs sense a broad spectrum of chemical signals, including hormones, ions, small molecules, neurotransmitters, and metabolites, and translate them into intracellular responses through finely tuned signal transduction pathways^3–6^. These responses are mediated either through orthosteric or allosteric ligand engagement with the cognate receptor^7^. Modulation of GPCR activation can profoundly alter cellular physiology^6,8,9^ and is implicated in numerous pathological conditions, including oncological, metabolic, cardiovascular, and central nervous system disorders^10–13^. Consequently, GPCRs are targets of approximately one-third of all approved therapeutic drugs5,^14,15^.

Despite their central biological and pharmacological importance, most GPCRs remain without selective therapeutic ligands^5^. One major challenge in targeting GPCRs is achieving high selectivity among closely related receptor subtypes^16^. Structural studies have established that GPCRs exist in dynamic equilibria between multiple active and inactive conformational states, further complicating ligand development^17^. Although small molecules dominate the GPCR drug landscape, innovative strategies to identify synthetic ligands, particularly nucleic acid–based ligands, that recognize GPCRs in their native cellular context remain limited. Aptamers are short, single-stranded nucleic acids (DNA, RNA, XNA, or the incorporation of unnatural base pairs)^18–20^ that fold into defined three-dimensional structures capable of binding to targets with high affinity and specificity^21–24^. They are generated through Systematic Evolution of Ligands by Exponential Enrichment (SELEX), a combinatorial selection process that enables the discovery of nucleic acid ligands against purified proteins, membrane receptors, and whole cells^25,26^. However, we previously introduced an advanced selection platform, termed as Ligand-Guided Selection (LIGS), that enables the discovery of highly selective aptamers against complex receptors in their native state^27,28^. Taking advantage of the inherent competition between weak and strong binders, LIGS uses a secondary ligand, such as an antibody with high affinity towards the targeting receptor, to selectively elute receptor-bound aptamers from an enriched library^29^. To the best of our knowledge, the use of small-molecule ligands to drive selective elution and exploit receptor conformational states has yet to be explored.

Here, we expanded LIGS to exploit small-molecule, ligand-induced conformational switching of GPCRs from inactive to active state to discover high-affinity, highly specific DNA aptamers targeting the β_2_-adrenergic receptor (β_2_AR) via LIGS using overexpressing HEK293 (W9) cells. β_2_AR, a prototypical class A GPCR, has been extensively studied and serves as a model for understanding GPCR activation mechanisms^7,30,31^. We applied a modified LIGS platform with four ligands, consisting of two agonists (isoproterenol and epinephrine), one antagonist (propranolol), and a FLAG monoclonal antibody, to selectively elute aptamers bound to β_2_AR from an enriched cell-SELEX library against W9 cells. Enriched Cell-SELEX and LIGS libraries were then subjected to high-throughput sequencing, and the resulting sequences were analyzed using the FASTAptamer toolkit. We then used a previously introduced LIGS-specific Galaxy bioinformatics platform to identify specific sequence families^32,33^. Multiple sequence alignment using ClustalW^34^ and the Galaxy platform identified twenty families from which candidate aptamers were selected based on fold enrichment and condition-specific representation across LIGS conditions. Three high-affinity DNA aptamers specific to β_2_AR in its native state were subsequently identified and validated. All three aptamers competitively bind β_2_AR with high specificity and selectively internalize into β_2_AR-positive cells. Importantly, their reduced affinity for purified receptor relative to cell-surface β_2_AR suggests preferential recognition of receptor context stabilized within the native membrane environment, consistent with conformational selectivity. Collectively, this work demonstrates that DNA aptamers can be evolved to selectively target a G protein– coupled receptor in its native cellular state. By harnessing ligand-induced conformational switching during selection and high throughput data analysis, we establish ligand-guided conformational selection as a broadly applicable DNA aptamer elution strategy for discovering nucleic acid ligands against GPCRs in their native state.

## RESULTS

### Cell-SELEX-based enrichment of Aptamers for β_2_AR recognition

To facilitate DNA aptamer development against β_2_AR, W9 cells overexpressing β_2_AR were used for aptamer selection. Receptor expression on the W9 cells was verified by anti-FLAG immunostaining targeting the engineered N-terminal FLAG tag using flow cytometry **(Figure S1A-C)**. SELEX against W9 cells was initiated with a diverse ssDNA library and progressively refined over multiple selection rounds under increasingly stringent conditions **(Table S1)**. Selection dynamics were monitored by high-throughput sequencing (NovaSeq 6000) and analyzed using FASTAptamer. The starting library (round 0) represented maximal sequence diversity, whereas successive rounds showed progressive enrichment of specific sequences **(Figure 1A and B)**, consistent with evolutionary selection and amplification of higher affinity β_2_AR binders. To eliminate nonspecific binders, counter-selection was introduced using HeLa cells, which express minimal β_2_AR compared to wild-type HEK293T cells, as confirmed by mRNA analysis **(Figure S2A and B)**. Three additional rounds of negative selection were performed to deplete sequences binding to nontarget surface molecules. Subsequent selection rounds were sequenced to monitor library evolution, confirming progressive enrichment of dominant sequences **(Table S2)**. To monitor selection progression, enriched aptamer pools from rounds 2, 4, 7, 9, 11, and 14 were analyzed via flow cytometry. A progressive increase in fluorescence intensity upon binding to W9 cells was observed across these rounds relative to the fluorescently labeled round 0 library control **(Figures 1C and S3)**. This rightward shift and observed progressive sequencing-based enrichment analysis indicates accumulation of specific binders towards W9 cells.

**Figure 1.**
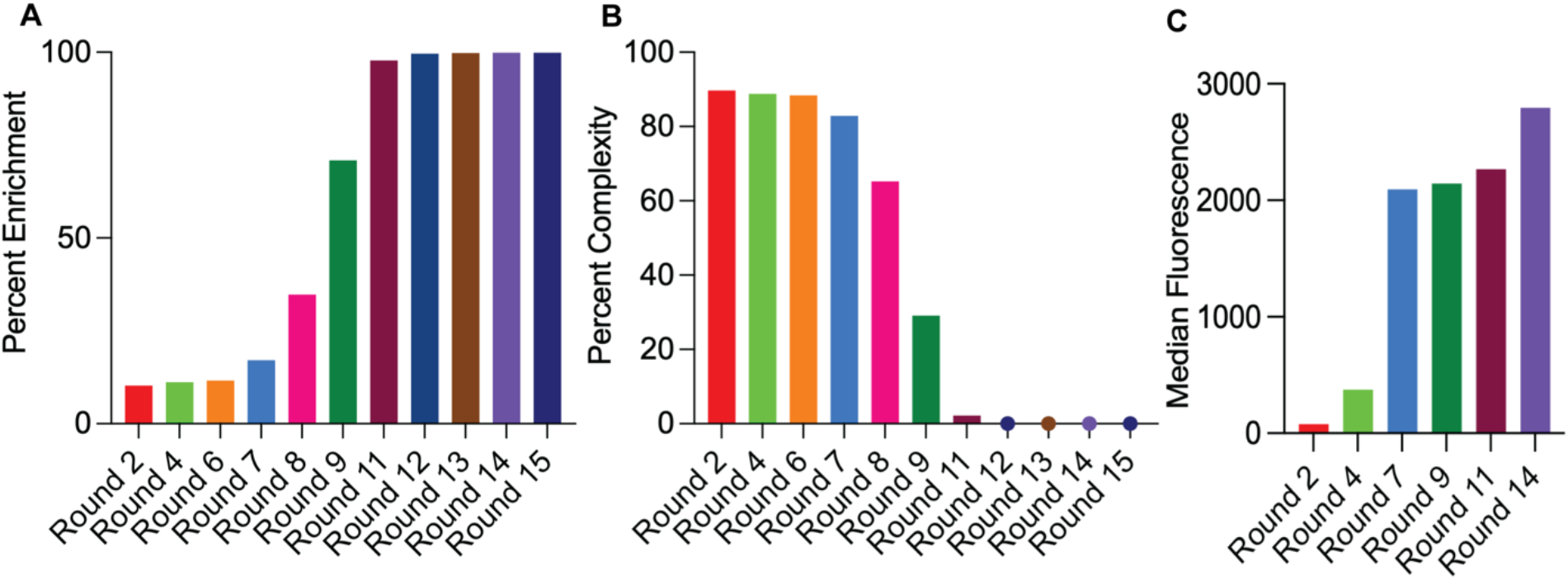
Cell-SELEX enrichment of library. **A-B)** The enrichment and complexity of cell-SELEX libraries were monitored by high-throughput sequencing. The percentage of enrichment across the sequenced rounds of cell-SELEX was calculated as percent enrichment = [(1 - complexity) x 100]. **C)** Progress of the selection was assessed by incubating 1.5 × 10^5^ W9 cells with 12.5 pmol of the fluorophore-labeled libraries from various rounds of cell-SELEX. The changes in median fluorescence intensity were quantified through flow cytometry. No shift in fluorescence intensity was observed in rounds 2-4 compared with the fluorophore-labeled round zero library. A gradual increase in fluorescence intensity was observed from round 4 to 14.

### Isolation of β_2_AR Aptamers via Ligand-Guided Selection (LIGS)

The ligand-guided selection (LIGS) strategy for the isolation of β_2_AR aptamers is outlined in Scheme 1, and ligands are summarized in Table 1. Prior to LIGS, the apparent affinities of enriched libraries from rounds 11 and 14 were evaluated against W9 cells. Round 11 library exhibited an apparent *K*_*d*_ of 41.1 nM, whereas round 14 library showed a *K*_*d*_ of 60.5 nM under conditions of total library binding **(Figure S4)**, indicating substantial enrichment of the Cell-SELEX library towards W9 cells. We next determined the affinities of the secondary ligands used to drive selective elution. The FLAG monoclonal antibody displayed a *K*_*d*_ of 2.0 nM **(Figure S5D)**, while epinephrine, isoproterenol, and propranolol displayed EC_50_/IC_50_ values of 21.4 nM, 12.1 nM, and 198.2 nM, respectively **(Figure S5A–C)**. These affinity measurements informed the design of LIGS conditions to competitively and selectively elute β_2_AR-specific aptamers. An isotype control was incorporated to discriminate between high off-rate and off-target binders **(Table 1)**. Libraries generated under each LIGS condition were subsequently subjected to next-generation sequencing and analyzed using FASTAptamer and the Galaxy platform, as previously described^*32*^.

**Table 1.**
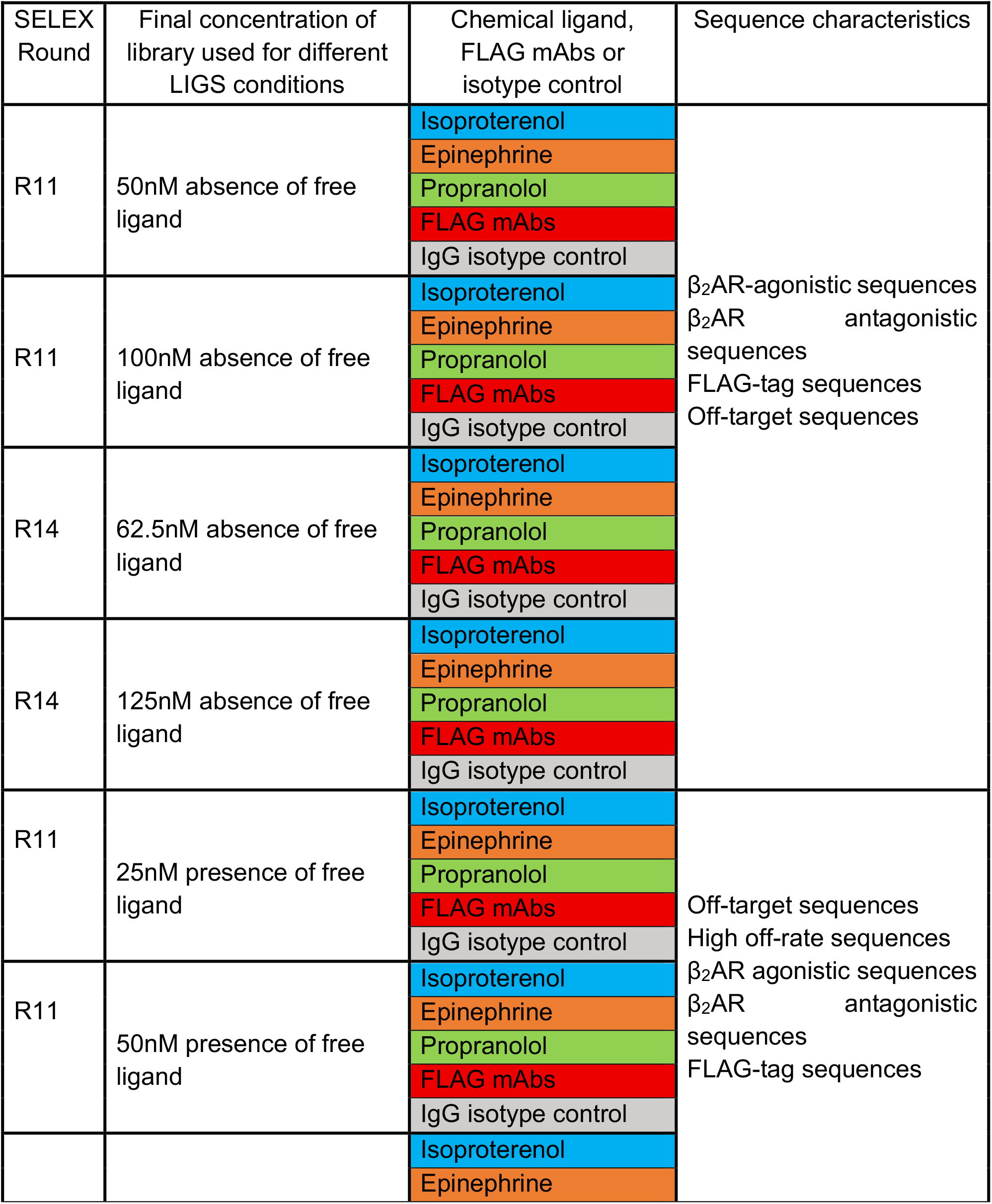
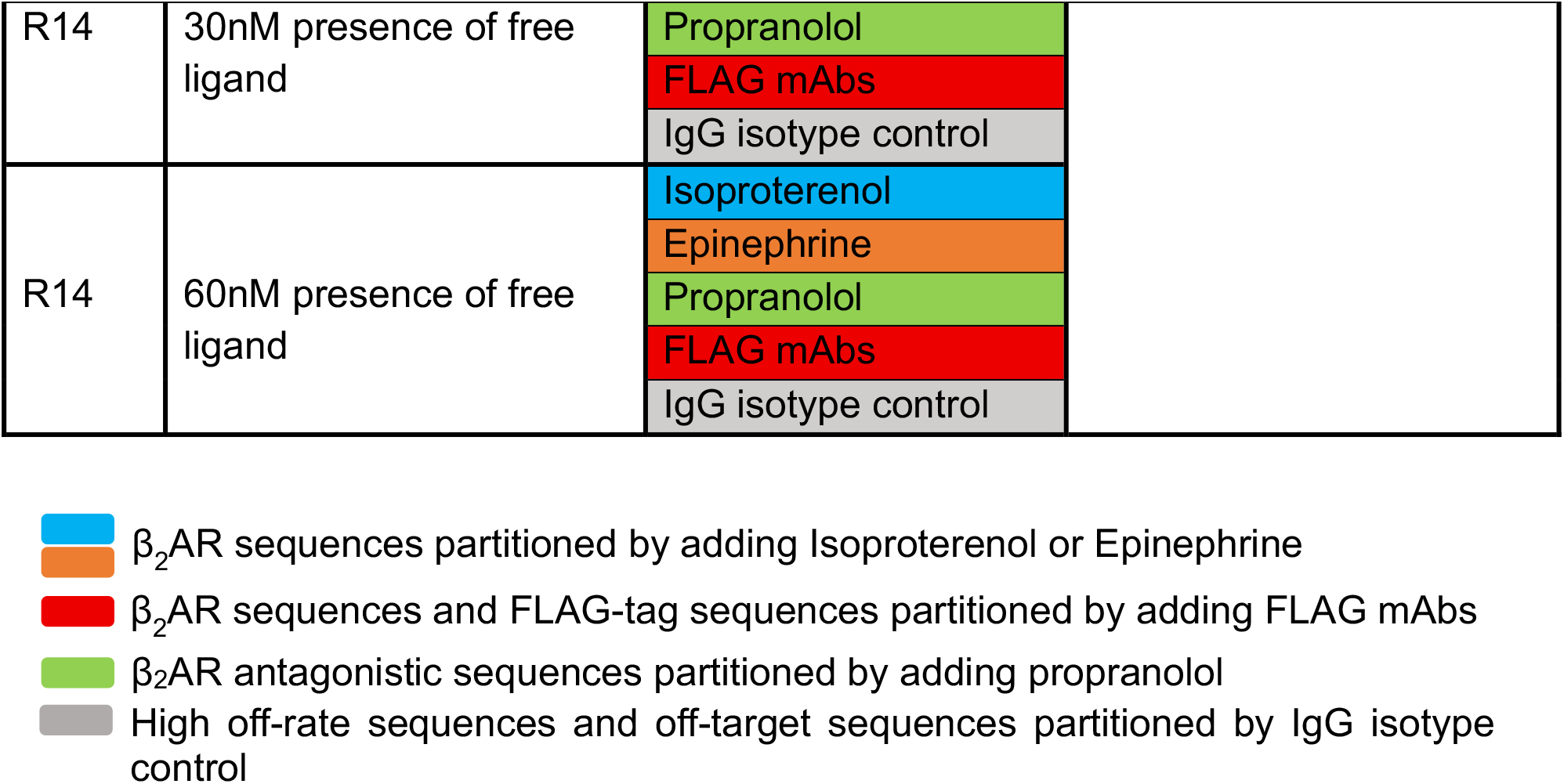
Ligand-guided selection conditions utilized to outcompete sequences bound to the β-2 adrenergic receptor and predicted characteristics of the outcompeted sequences.

**Scheme 1.**
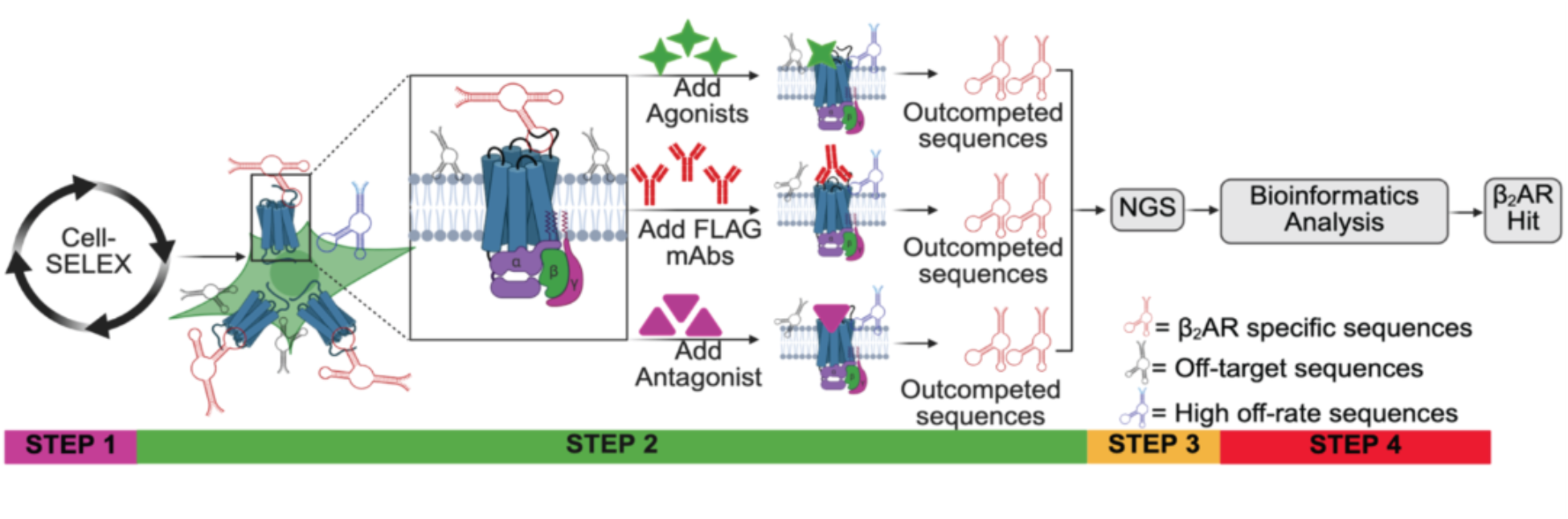
Ligand-guided selection and identification of β_2_AR-specific DNA aptamers. The schematic outlines the integrated experimental and bioinformatics workflow used to isolate β_2_AR-targeting aptamers. **Step 1**. Fifteen rounds of whole-cell SELEX selection were performed on β_2_AR-overexpressing W9 cells, including three counterselection rounds against HeLa cells to eliminate nonspecific binders. **Step 2**. Ligand-guided selection (LIGS) of β2AR-bound sequences using excess epinephrine, isoproterenol, propranolol, or FLAG monoclonal antibody to exploit ligand-induced receptor conformations as a strategy to elute specific aptamers. **Step 3**. Eluted libraries derived from enriched rounds (R11 and R14) were subjected to high-throughput sequencing (NovaSeq 6000). **Step 4**. Sequencing datasets were analyzed using FASTAptamer and the Galaxy platform, applying iterative enrichment and comparative filtering to discriminate β_2_AR-specific binders. Multiple sequence alignment identified distinct sequence families from which representative candidates were selected for downstream biochemical vslidation for affinity and specificity.

### Bioinformatics analysis-driven identification of high-priority β_2_AR aptamer candidates

Following adapter trimming, sequences were ranked by abundance (reads per million (RPM)) using FASTAptamer-count **(Figures S6 and S7)**. Fold enrichment under each LIGS condition was calculated with FASTAptamer-enrich (**Table S2**) according to established parameters. Enrichment ratios were defined as follows: RPMy/RPMx (isotype-eluted sequences relative to enriched SELEX pools), RPMz/RPMx (β_2_AR ligand–eluted sequences relative to enriched SELEX pools), and RPMz/RPMy (β_2_AR ligand–eluted sequences relative to isotype-eluted sequences), where x denotes the enriched SELEX round, y is the isotype control condition, and z is the β_2_AR-specific ligand condition. Downstream analysis was performed using the Galaxy platform. For each ligand (epinephrine, isoproterenol, propranolol, and FLAG mAb), enrichment values (z/x) were plotted against sequence abundance to define condition-specific thresholds **(Figures S8 and S9)**. A cutoff of ≥2-fold enrichment was applied to identify sequences selectively enriched under β_2_AR ligand competition relative to enriched SELEX pools. Application of the initial z/x ≥2 filter retained approximately 0.28–4.79% of sequences across ligand conditions **(Table S3)**. To further refine specificity, a second threshold (z/y ≥5; y = isotype control) was applied to the filtered pools **(Figures S10 and S11)**, retaining 3.13–23.94% of sequences **(Table S3)**. A subsequent cross-comparison step was implemented to distinguish ligand-specific binders from nonspecific or membrane-disruption artifacts. Sequences enriched in agonist-(isoproterenol, epinephrine) or FLAG mAb–eluted pools were compared against propranolol-eluted pools, and vice versa, retaining only non-overlapping sequences **(Figure 2A)**. To eliminate sequencing artifacts, sequences supported by fewer than two reads were excluded. The remaining sequences were ranked by abundance, converted to FASTA format **(Figure 2A)**, and subjected to multiple sequence alignment (ClustalW) **(Figure 2A)**. This analysis identified 20 distinct sequence families. Based on fold enrichment and ligand-specific occurrence, one representative sequence from each family was selected as a potential hit **(Table S4)**. Evolutionary trajectories of these candidates were subsequently tracked across sequenced SELEX rounds **(Figure 2B)**.

**Figure 2.**
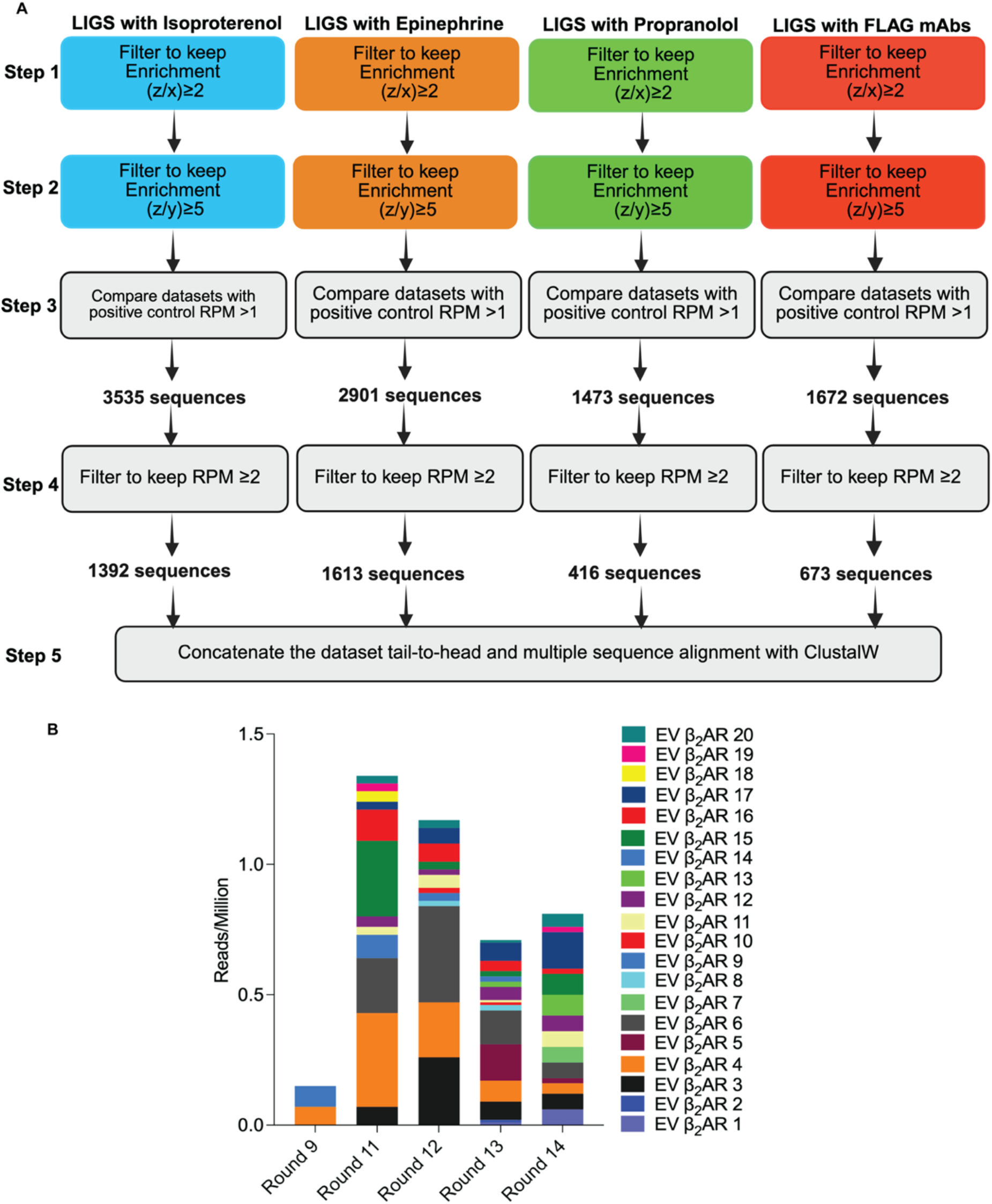
Bioinformatics Analysis of HT Sequencing Data from LIGS Libraries. **A)** Schematic diagram summarizing downstream analyses of FASTAptamer-Enrich data from different LIGS conditions with the R11- and R14-enriched SELEX libraries using the GALAXY bioinformatics toolkits. Filtering steps were employed as outlined to discriminate potential β_2_AR-specific binders, showing the total number of sequences obtained for each ligand. **B)** Evolution of β_2_AR hit sequences in different rounds of cell-SELEX.

### Validation and affinity determination of specific aptamer candidates

β_2_AR hit candidates identified through LIGS and bioinformatics analyses were evaluated for their binding affinity to W9 (β_2_AR-overexpressing) and HeLa (counterselection) cells. Whole-cell fluorescence assays using FAM-labeled aptamers at 4 °C revealed that 15 of 20 candidates exhibited >4-fold higher binding to W9 cells relative to that of HeLa cells **(Figure S12A and Figure S13)**. Binding specificity was assessed by comparing median fluorescence intensity to that of a randomized control sequence, indicating selective recognition of a target enriched on W9 cells. Apparent binding affinities were determined by incubating W9 cells with increasing concentrations (5–250 nM) of fluorophore-labeled candidates under equilibrium conditions. Nonlinear regression of fluorescence signals, normalized to background, yielded saturable binding curves consistent with receptor engagement. All fifteen candidates bound in the nanomolar range **(Figure S12B and S14)**. EV β_2_AR 19 exhibited the highest apparent affinity (*K*_*d*_ = 84.0 nM; Figure 3), whereas EV β_2_AR 13 displayed the lowest affinity (*K*_*d*_ = 780 nM). These results confirm that the selected aptamers bound β_2_AR with high affinity and specificity in the native cellular context.

**Figure 3.**
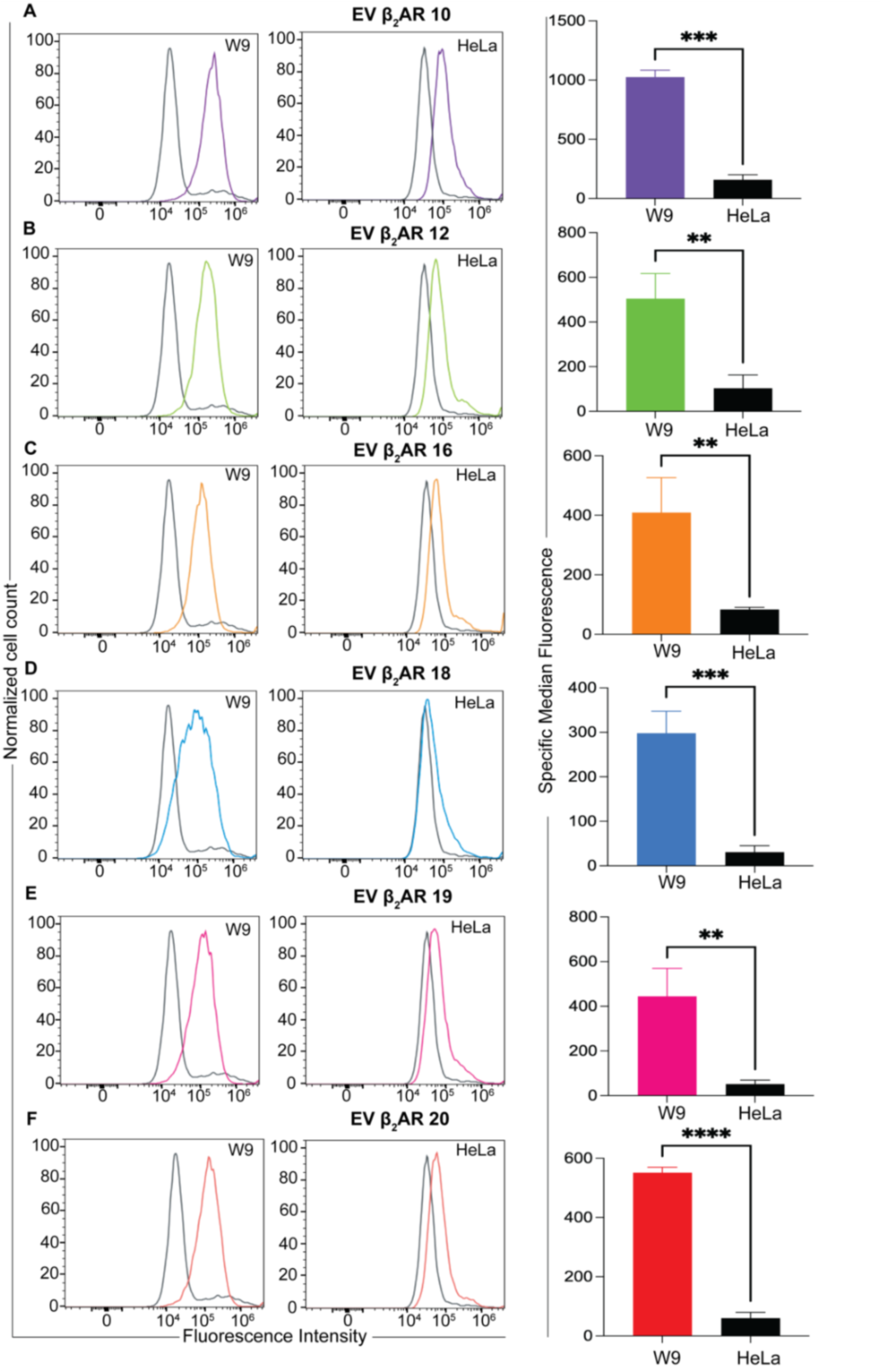
The specificity of hit aptamer candidates for binding to W9 cells. **A-F)** Binding of hit aptamer candidates to W9 cells was examined by incubating 7.5 × 10^4^ W9 or HeLa cells with 250 nM of fluorescently labeled hit aptamer candidate or a random sequence at 25 °C for 45 minutes. Subsequently, binding was quantified through flow cytometry, and specific median fluorescence (Median Fluorescence Intensity, MFI) was calculated as MFI = [median fluorescence of aptamer candidate − median fluorescence of random control sequence]. Fluorescence intensity measurements demonstrate strong binding to W9 cells overexpressing β_2_AR, but only minimal signal detection in HeLa cells, indicative of β_2_AR specificity on W9 cells. The specific median fluorescence of each candidate was plotted using GraphPad Prism software and further analyzed via a non-parametric *t*-test (*p*-values ranging from <0.0001 to 0.0087), and data represent the mean ± SD of three independent experiments.

### Assessment of β_2_AR Specificity

Specificity of the β_2_AR hit candidates was evaluated by comparing binding to W9 (β2AR-overexpressing) and HeLa cells. Whole-cell fluorescence assays at 4 °C demonstrated robust and statistically significant binding of all fifteen candidates to W9 cells relative to that of HeLa controls **(Figure S15)**, confirming selective recognition of β_2_AR-enriched cells. Based on apparent affinity, six candidates (EV β_2_AR 10, 12, 16, 18, 19, and 20) were selected for further evaluation at 25 and 37 °C. All six retained statistically significant preferential binding to W9 cells under these conditions **(Figure 3 and S16)**. To further assess target specificity, W9 cells were pretreated with 10 μM isoproterenol to induce β_2_AR internalization prior to aptamer binding analysis. As expected, isoproterenol significantly reduced surface FLAG antibody binding, confirming receptor internalization. Similarly, three candidates, EV β_2_AR 12, 16, and 19, also showed a significant decrease in binding following agonist treatment **(Figure 4)**, consistent with β_2_AR-dependent recognition. Although EV β_2_AR 10, 18, and 20 exhibited reduced binding after agonist treatment, these changes did not reach statistical significance. Collectively, these results identify EV β_2_AR 12, 16, and 19 as β_2_AR-specific aptamers that recognize receptor populations present at the cell surface and respond to ligand-induced receptor internalization.

**Figure 4.**
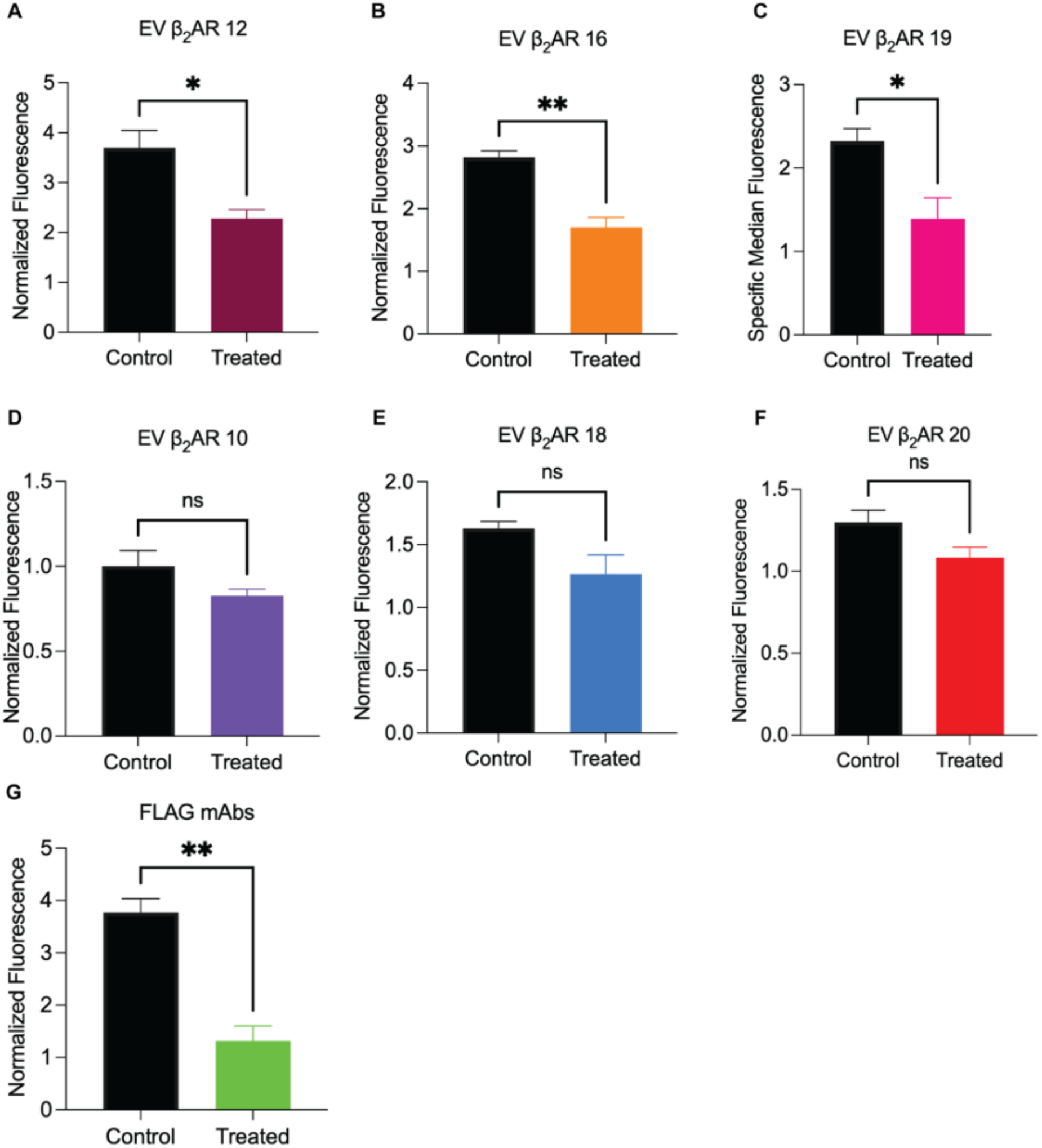
Specific binding of hit aptamer candidates to the beta-2 adrenergic receptor. W9 cells were pretreated with 10 μM of isoproterenol at 37°C for 45 minutes. Following treatment, the pretreated cells were stained with A) FLAG mAbs and B-G) FAM-labeled hit β_2_AR aptamer candidates. Binding to β_2_AR was quantified by measuring the fluorescence intensities with flow cytometry. Specific median fluorescence (Median Fluorescence Intensity, MFI) was calculated as MFI = [median fluorescence of aptamer candidate − median fluorescence of random]. A decrease in binding is indicative of the depletion of β_2_AR from the cell surface, which results from internalization induced by isoproterenol. The specific median fluorescence of each candidate was plotted using GraphPad Prism software and further analyzed via a non-parametric *t-*test (*p* = 0.0030, 0.0070, and 0.0455 for EV β_2_AR 12, 16, and 19, respectively); all experiments were performed in triplicate.

### Aptamers are highly specific towards native β_2_AR

To determine whether the identified β_2_AR aptamers recognize an epitope overlapping, or proximal to, the FLAG-tag region, we performed whole-cell competition assays using β_2_AR-overexpressing W9 cells. Cells were preincubated with either CY3-labeled EV β_2_AR 16 or FLAG monoclonal antibodies, followed by assessment of FAM-labeled aptamer binding by flow cytometry. Pretreatment significantly reduced binding of EV β_2_AR 12, 16, and 19 compared to untreated controls **(Figure 5A-F)**, indicating competitive interference. Next, we used microscale thermophoresis (MST) to characterize the binding of aptamers EV β_2_AR 16 and EV β_2_AR 19 to purified β_2_AR, using BSA as a control^35,36^. Cy5-labeled aptamers were used as the fluorescent ligand, and purified β_2_AR served as the unlabeled target. Both aptamers exhibited greater binding to β_2_AR than binding to BSA **(Figure 5G-H)**, with EV β_2_AR 19 showing modestly higher affinity than that of EV β_2_AR 16. To further evaluate specificity, we compared binding of EV β_2_AR 16 and EV β_2_AR 19 to purified β_2_AR against a randomized sequence **(Figure S18)**. As expected, the randomized aptamer showed minimal interaction with β_2_AR, displaying a markedly reduced binding response relative to the selected aptamers. Consistent with cell-based measurements, EV β_2_AR 16 and EV β_2_AR 19 exhibit high-affinity binding to W9 cells. In contrast, binding to purified β_2_AR required higher concentrations of both aptamer and protein. Additionally, occupation of the receptor by CY3-labeled EV β_2_AR 16 diminished subsequent binding of FAM-labeled aptamers. These findings demonstrate that the three aptamers compete with one another and with FLAG monoclonal antibodies for receptor engagement, consistent with recognition of overlapping, or spatially proximal, epitopes near the N-terminus.

**Figure 5.**
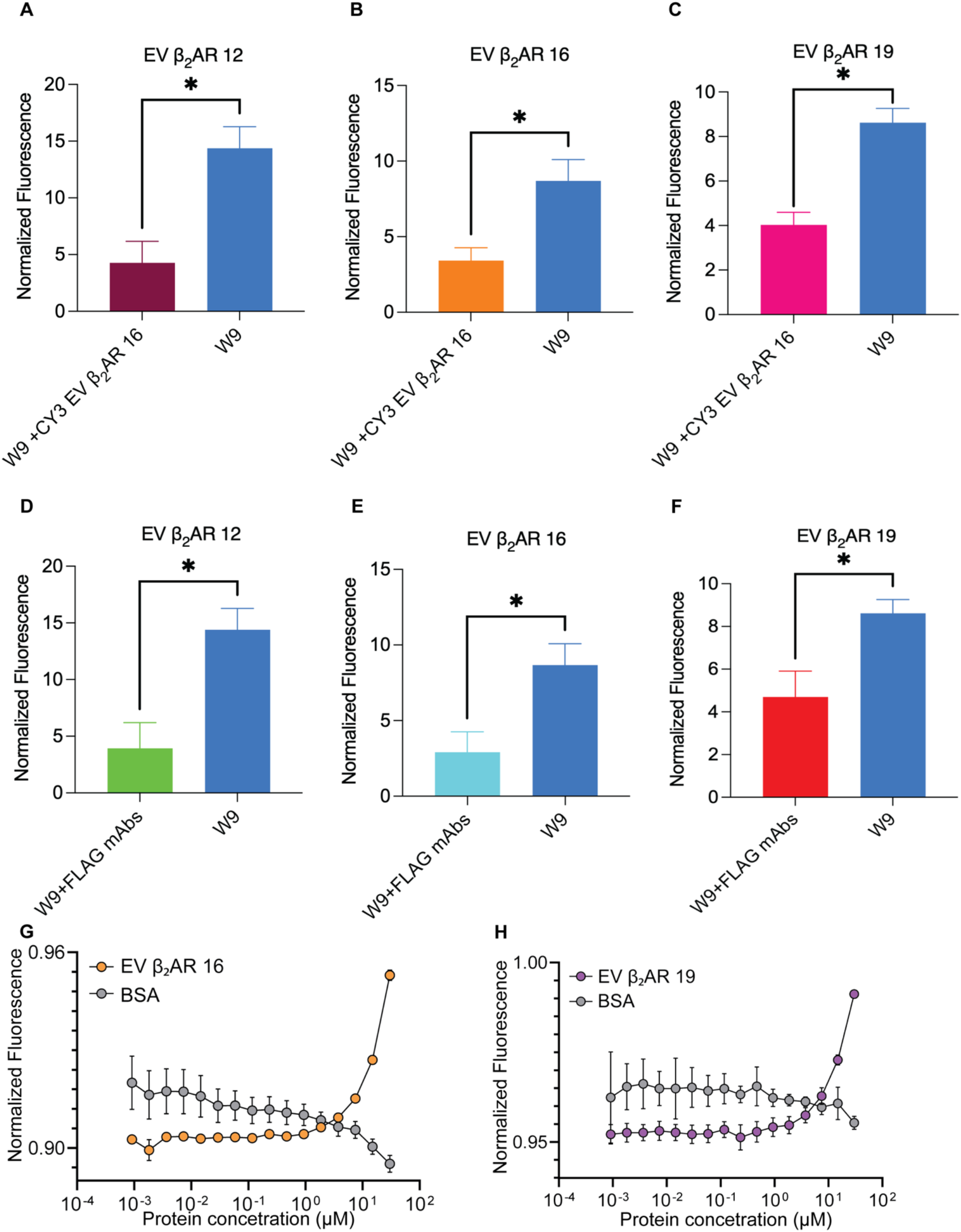
Specificity of aptamer towards the beta-2 adrenergic receptor. **A-F)** Fluorescence-based ligand competition assay in whole cells showing inhibitory binding of FAM-labeled aptamers in the presence of CY3-labeled EV β_2_AR 16 or FLAG mAbs. Fluorescence intensity was quantified through flow cytometry, and median fluorescence was normalized to signals obtained from the binding of FAM-labeled aptamers to random (serving as the control). The normalized median fluorescence values were calculated and graphically represented using GraphPad Prism. Bars depict the mean ± SEM across three independent experiments, and statistical significance was assessed employing one-way ANOVA. Asterisks denote statistically significant differences compared to the untreated control cells (*p* < 0.05). **G-H)** Specificity of aptamer binding with purified β_2_AR protein. A representative MST normalized fluorescence of Cy5-labeled aptamers in the presence of unlabeled purified β_2_-adrenergic receptor. A fixed aptamer concentration of 1 µM was titrated with increasing concentrations of the purified protein (9.1 nM to 30 µM). The X-axis shows increasing protein concentration, and the Y-axis shows normalized fluorescence measured on the Monolith X instrument. Data represented are mean ± SEM from three independent experiments. Statistical significance was determined using XY linear regression analysis (*p*-value <0.0001).

### Aptamer-induced internalization of β_2_AR

The three β_*2*_AR-specific aptamers were isolated from LIGS pools competitively eluted with two agonists, epinephrine and isoproterenol. Given that these agonists promote β_*2*_AR internalization, we examined whether the aptamers could similarly induce receptor trafficking. To accomplish this, W9 cells were incubated with FAM-labeled aptamers at 37 °C, and binding was monitored over time by flow cytometry. Median fluorescence intensity increased progressively from 0 to 60 minutes **(Figure 6A-C)**, reflecting cumulative surface binding and receptor internalization. To distinguish internalized receptors from surface-bound complexes, cells were treated with aptamers for 60 minutes and subsequently trypsinized to remove surface receptors, leaving internalized aptamer–receptor complexes intact. Trypsinized samples were compared with non-trypsinized controls and a randomized sequence control. Flow cytometric analysis revealed substantial receptor internalization induced by EV β_*2*_AR 12 and 19 with approximately 80–90% of bound aptamers internalized after 60 minutes. EV β_*2*_AR 16 also promoted internalization, albeit to a lesser extent (approximately 40–60%) **(Figure 6A–C)**. Confocal microscopy confirmed intracellular redistribution of the aptamer–receptor complexes **(Figure 6A–C)**. These findings indicate that agonist-eluted aptamers not only bind β_*2*_AR but also promote receptor internalization, consistent with the functional engagement of native receptor conformations.

**Figure 6.**
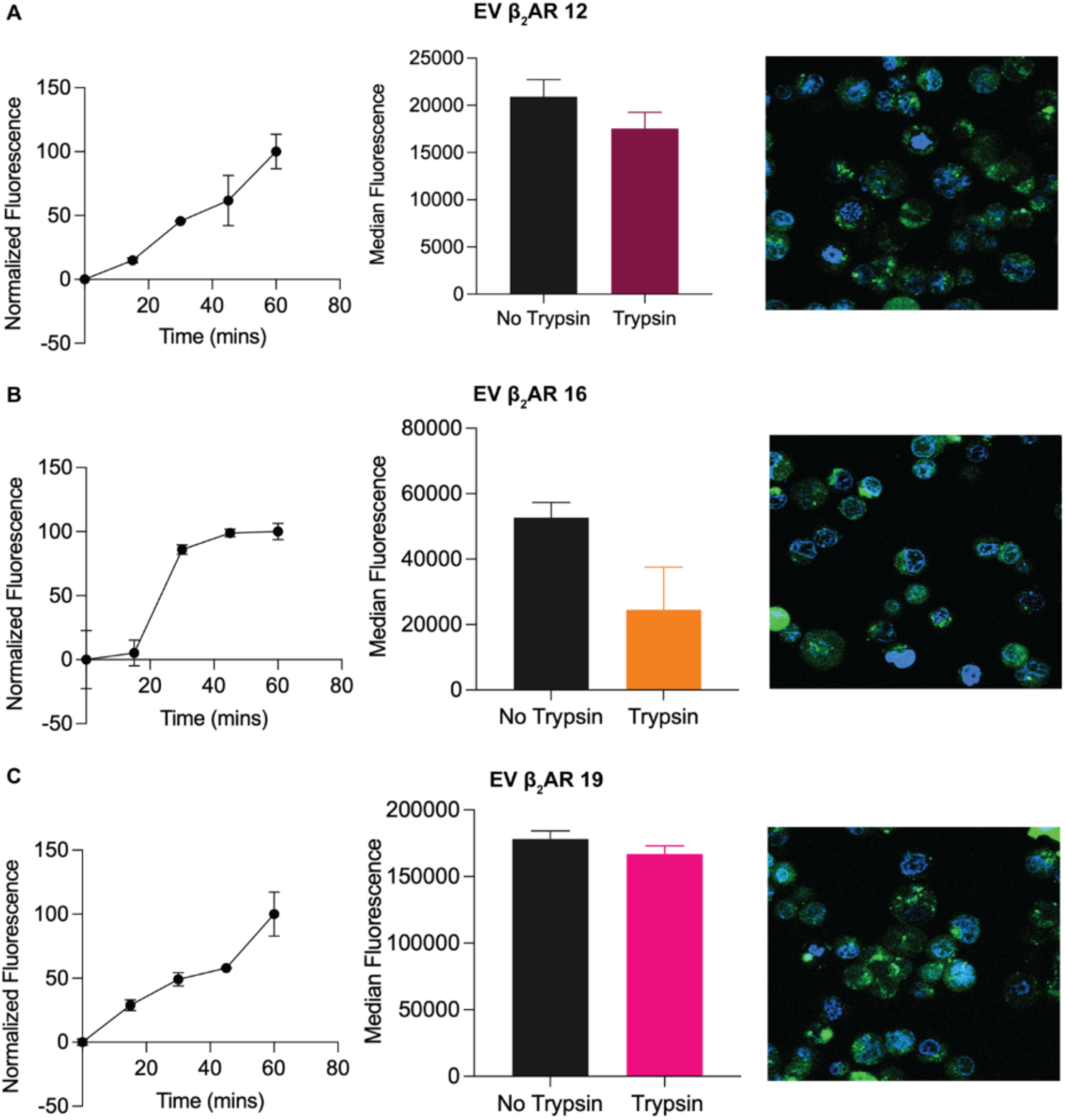
Aptamers mediate internalization of β_*2*_AR. **A-C)** W9 cells were exposed to 250 nM of aptamers for various durations (0 to 60 minutes) at 37°C, followed by measurement of fluorescence via flow cytometry. The median fluorescence values were plotted against time and analyzed with GraphPad Pr using XY analysis. An increase in fluorescence indicates total binding (internal and external) of the aptamers to the cells. W9 cells were incubated with 250 nM of aptamers for 60 minutes at 37°C. Following incubation, one group of cells was trypsinized to remove surface receptors, while the control group was left untrypsinized. The change in fluorescence intensity was measured by flow cytometry. The median fluorescence was plotted and analyzed with GraphPad Prism. The difference in fluorescence between the control and trypsinized groups reflects the proportion of aptamer-bound β_2_AR that was internalized. Confocal microscopy images of trypsinized cells depict the intracellular localization of FAM-labeled aptamers (green) and the cell nucleus (blue).

## Discussion

Responding to a wide range of environmental cues, G protein–coupled receptors (GPCRs) constitute one of the largest families of membrane proteins^3^. These receptors function at the interface between extracellular and intracellular environments and transduce signals through downstream second-messenger pathways. Based on their central biological roles, approximately one-third of currently approved drugs target GPCRs^5^. Among GPCRs, β-adrenergic receptors play critical roles in mediating catecholamine-driven physiological responses^37^. The β2-adrenergic receptor (β2AR), in particular, is broadly expressed across multiple tissues and regulates such processes as airway smooth muscle relaxation and metabolic signaling through activation by epinephrine and norepinephrine^38,39^. Synthetic small-molecule antagonists (β-blockers) and selective agonists have been widely developed to modulate β2AR activity^40,41^. However, conventional small-molecule ligands primarily target conserved orthosteric sites, limiting subtype selectivity and constraining the ability to precisely control receptor conformational states^42^. These limitations highlight the need for programmable molecular ligands capable of targeting GPCRs with improved structural and functional specificity.

Nucleic acid ligands, such as aptamers, provide molecules for developing programmable molecular tools against membrane receptors. Aptamers can adopt complex three-dimensional structures that enable high specificity and affinity toward diverse biological targets, and they have been widely applied in diagnostics, molecular tool development, and therapeutic delivery^43^. Advances in chemically modified and unnatural nucleic acids have further expanded the structural and functional diversity available for ligand engineering^44–46^. Despite these advantages, generating aptamers against GPCRs remains challenging because receptors purified in detergent environments often lose native conformational features required for accurate ligand recognition. To address this limitation, we previously developed Ligand-Guided Selection (LIGS), a strategy that leverages competitive displacement within enriched SELEX libraries to selectively isolate receptor-binding aptamers directly from cells expressing membrane-bound targets^27,29,32^. Using this approach, receptor-specific aptamers have been generated against multiple immune receptors, including IgM, CD3, CD19, and CD20^27,29,32^. Building on this framework, we herein extended LIGS to target a structurally dynamic GPCR, β_2_AR, in its native membrane environment.

In this study, W9 cells engineered to overexpress β2AR were used to enable selection under physiologically relevant conditions. Four β2AR-directed ligands were incorporated into LIGS to selectively elute receptor-bound sequences: two agonists (isoproterenol and epinephrine), one antagonist (propranolol), and a FLAG monoclonal antibody recognizing the engineered extracellular epitope. Because these ligands stabilize distinct receptor conformations, this strategy enabled enrichment of aptamers capable of recognizing conformationally defined receptor states. High-throughput sequencing followed by bioinformatics analysis using FASTAptamer and Galaxy identified three specific aptamers (EV β_2_AR 12, 16, and 19), all of which were preferentially eluted under agonist competition conditions, suggesting conformational sensitivity toward agonist-stabilized receptor states.

Achieving subtype specificity for β_2_AR is inherently challenging owing to strong structural similarity among adrenergic receptor subtypes^47^. Nevertheless, all three aptamers demonstrated strong binding to β_2_AR-overexpressing W9 cells, while exhibiting minimal binding to low-β_2_AR-expressing HeLa cells, thus confirming receptor-dependent specificity. Functional validation through receptor internalization experiments further supported surface-specific recognition since agonist-induced internalization reduced aptamer binding in parallel with FLAG antibody controls. Competition assays revealed that the three aptamers compete, suggesting overlapping, or spatially proximal, binding regions, consistent with previous observations that LIGS-derived aptamers can converge on accessible extracellular domains.

Interestingly, aptamer binding affinity was significantly reduced when purified β_2_AR was used instead of membrane-displayed receptor, as determined by MST measurements which showed approximately five-fold lower affinity toward purified protein. Similar behavior was previously observed for membrane-targeted anti-IgM aptamers. Both cases support a key conceptual feature of the LIGS strategy: that aptamers preferentially recognize membrane-dependent structural or conformational states not fully preserved in purified systems. These findings reinforce the importance of native membrane context for ligand discovery against dynamic receptor systems such as GPCRs. Additionally, these findings highlight the advantage of ligand-guided selection (LIGS), which enriches aptamers against targets in their native cellular context. Beyond receptor recognition, we further demonstrate that the selected β_2_AR aptamers induce rapid receptor internalization. This observation suggests that the aptamers may stabilize receptor conformations that favor β-arrestin recruitment or may promote steric clustering that triggers endocytic pathways distinct from classical small-molecule activation mechanisms^48–51^. The ability of aptamers to modulate receptor trafficking introduces new opportunities for developing functionally selective GPCR ligands, as well as targeted drug delivery.

For the first time, this work collectively establishes LIGS-enabled discovery of conformationally sensitive aptamers targeting GPCRs in their native membrane environments, while simultaneously revealing functional modulation of receptor behavior. Compared with antibody-based approaches, aptamers offer structural programmability, tunable chemical modification, and flexible pharmacokinetic optimization, providing a versatile platform for GPCR-targeted therapeutic and chemical biology applications.

## Acknowledgements

PM gratefully acknowledge support from NIGMS grant R35GM139336, and the G-RISE fellowship awarded T32GM136499 to YNC, U-RISE fellowship 5134GM149459 to DS. AG and MB acknowledge support from R35 GM133589.

## Materials and Methods

### Cell culture and reagents

β_2_AR-overexpressing HEK293 cells (W9 cells) were a generous gift from the Lefkowitz lab at Duke University, North Carolina. HEK 293T cells endogenously expressing β_2_AR and HeLa CCL-2 cells were purchased from the American Type Culture Collection (ATCC, Manassas, VA, USA). W9, HEK 293T, and HeLa cells were maintained in high glucose DMEM and EMEM medium, respectively, supplemented with 10% dialyzed fetal bovine serum, 100 units/mL (10,000IU penicillin and 10,000μg/mL streptomycin, Corning), and 1% MEM non-essential amino acids at 37°C in a 5% CO_2_ incubator. Flow cytometry was regularly used to monitor β_2_AR-expressing W9 cells using monoclonal antibodies (Rabbit anti-human monoclonal, R&D Systems) against the FLAG tag.

### Chemical ligands and antibodies

The β_2_AR agonists (epinephrine, isoproterenol) and antagonist (propranolol) were purchased from Sigma Aldrich. Rabbit IgG mAb (cat#: IC8529G) against the FLAG epitope and Isotype control Rabbit IgG mAb (cat#: IC1051G) were purchased from R&D Systems.

### SELEX and cell suspension buffers

The SELEX buffer (Tris-buffered saline; TBS) was formulated with TRIS-HCL in nuclease-free water and composed of 20mM TRIS-base/HCl, 145mM NaCl, 5.4mM KCl, 0.8mM MgCl2, 1.8mM CaCl_2_ with pH 7.4. To block nonspecific binding sites on the cell surface and culture plate, the cell suspension buffer (derived from the SELEX buffer) was supplemented with 200 mg/L tRNA (Sigma-Aldrich), 2g/L bovine serum albumin, and 200 mg/L salmon sperm DNA.

### SELEX library and primers

DNA library and primers were adapted from Liu et al. (2012) and consisted of a hybridization region (constant 20 bases in the 5’ and 3’ ends), flanking a 37-base randomized region (5’-ATG AGA GCG TCG GTG TGG TA-N_37_-T ACT TCC GCA CCC TCC TAC A-3’)^52^. The forward and reverse primers were 5’-labeled with Fluorescein dye (5’-6-FAM-ATG AGA GCG TCG GTG TGG TA) and biotin tag (5’-Biotin-T ACT TCC GCA CCC TCC TAC A), respectively. Both primers and DNA library were used to optimize cell-SELEX amplification conditions. PCR mixtures consisted of 0.25 units of Takara Hot Start Taq polymerase, 5 μM forward primer, 5 μM reverse primer, 2.5 mM dNTP mix, 10X PCR buffer, nuclease-free water, and 10 nM DNA template in a 25 μL total reaction volume. The amplification cycle started with an initial step at 98 °C for 3 mins, then denaturation at 98 °C for 30s, annealing at 60 °C for 30s, and elongation at 72 °C for 1min:30s, followed by a final extension at 72 °C for 15mins. The DNA library, primers, aptamers, and random controls were either synthesized and purified in-house using an ABI-394 synthesizer and reverse-phase HPLC or purchased from Integrated DNA Technologies Inc. (IDT).

### Cell-SELEX with adherent cells

W9 cells were cultured to approximately 90% confluency and washed three times with the SELEX buffer before incubation with the DNA library. For the first selection round, an initial concentration of 15 nmoles DNA library in 150 μL SELEX buffer was denatured at 95 °C for 10 mins and allowed to fold at 4 °C for 45 mins. The folded library’s volume was adjusted to 3ml with the cell suspension buffer before being added and incubated with the adhered cells for 45 minutes at 4 °C with intermittent shaking. After incubation, the cells were washed once with 9 mL of SELEX buffer and detached with 600 μL of SELEX buffer. Bound DNA was eluted by heating at 95°C for 15 minutes and then centrifuged at 4 °C for 15 minutes at 15,000 rpm. The eluent was first PCR amplified using five cycles to increase the number of copies of each sequence. The PCR amplicon was further subjected to 10 amplification cycles and used to regenerate the single-stranded DNA (ssDNA) library needed for the subsequent selection round. In the second selection round, template volume and cycle optimizations were performed using 10% and 20% of the DNA template in 25 μL of the PCR reaction with 8 and 10 cycles. The template volume was maintained at 10% of the total volume of the 25 μL PCR reaction for each selection round. The subsequent selection rounds were conducted as described above with minor modifications. The eluent from each selection round was used for cycle optimization, and the regenerated ssDNA was quantified using UV-vis spectroscopy, concentrated, and resuspended in the SELEX buffer to an initial concentration of 500 nM. Stringency in the selection process was applied starting from round 2. A progressive increase in the number of washes after incubation was performed, ranging from 3 mL x 2 at round 2 to 3 mL x 3 in subsequent rounds. Cell culture plate size was decreased from 100mm to 60mm at round 7. Three rounds of counterselection were performed with HeLa cells from round 13. Briefly, 300μL of the 500 nM library from round 12 were prepared as previously mentioned and used for positive selection. After positive selection, the cells were washed with 3mL x 3 of the SELEX buffer to remove the unbound sequences. The bound library was eluted and separated from the cells by centrifuging at 4°C for 15 minutes at 15000 rpm. The supernatant was collected for incubation with washed HeLa cells at 4°C for 45 minutes. After incubation, the supernatant with the unbound sequences was collected and PCR amplified. Amplicons were used for ssDNA regeneration for the subsequent round of selection.

### Monitoring cell-SELEX progress

The selection progress was monitored with flow cytometry analysis and high-throughput sequencing (Illumina sequencing). PCR amplification introduced a fluorescent tag (FAM) at the 5’-end of the DNA library used for flow cytometry. The DNA library (12.5 pmol in 25 μL) was prepared as in the SELEX procedure and incubated with 25 μL of washed W9 cells in suspension buffer for 45 minutes on ice. The incubated cells were washed and resuspended in 250 μL of the SELEX buffer. 10,000 events were analyzed with a Cytex NL-2000 flow cytometer. The DNA library from the selected rounds of cell-SELEX was prepared for Illumina sequencing and sequenced using NovaSeq 6000.

### Determination of apparent affinity of the enriched DNA library

The apparent affinity of the enriched SELEX library for the β_2_AR-overexpressing cells was determined using a range of library concentrations from 1 to 250 nM. Enriched libraries (round 11 or round 14) and control library (round zero) were prepared as previously mentioned, and each concentration was incubated with 1.5 × 10^5^ washed W9 cells in 25 μL of suspension buffer for 45 minutes on ice. After incubation, the cells were washed once with 1 mL of the SELEX buffer and resuspended in 250 μL of the SELEX buffer or resuspended without washing. Flow cytometry was used to record and analyze 10000 events. The apparent dissociation constant (*K*_*d*_) of the enriched library was determined from a plot of the specific median fluorescence of the enriched library against different concentrations, using nonlinear regression and site-specific binding analysis in GraphPad Prism.

### Ligand-guided selection after washing unbound primary ligands

Based on the apparent affinity of the enriched R11 and R14 rounds, two different final concentrations for each round were used in LIGS. The enriched library at round 11 (before counterselection) or round 14 (after counterselection) was used at a final concentration of 50 and 100 nM or 62.5 and 125 nM, respectively. The W9 cells were detached and washed three times with 3 mL of SELEX buffer. The washed cells were resuspended at a concentration of 1.5 × 10^5^ cells in 25 μL and incubated with 25μL of the doubled final concentration of the DNA library prepared as mentioned in the SELEX procedure for 45 minutes at 4°C. After incubation, the cells were washed once with 3 mL of the SELEX buffer and resuspended in 50 μL of CSB. The resuspended cells were incubated with 20 times the EC50, IC50, or *K*_*d*_ value of epinephrine or isoproterenol, propranolol and FLAG mAbs, respectively, or isotype control by adding 5μL of the respective ligand for 30 minutes at 4°C. After incubation, the cells were pelleted at 5000g for 1 minute, and the supernatant was collected and placed on ice for high-throughput sequencing preparation. Ligand-guided selection without washing unbound primary ligands

W9 cells were detached and washed, as previously described. The cells were resuspended at a concentration of 1.5 × 10^5^ cells in 25 μL and incubated with 25 μL of double the different final concentrations of round 11 (25 and 50 nM) or round 14 (30 and 60 nM) DNA library prepared as described in the SELEX procedure for 45 minutes at 4°C. After 45 minutes of incubation, 5 μL of ligand, Epinephrine, Isoproterenol, Propranolol, FLAG mAbs, or isotype control, were added to reach a final concentration 20 times the EC50, IC50, or *K*_*d*_ value for each ligand. The setup was incubated for 30 minutes at 4°C. After incubation, the cells were pelleted at 5000g for 1 minute, and the supernatant was collected and prepared for high-throughput sequencing.

### High-throughput Illumina sequencing sample preparation

Eluted DNA sequences from the different LIGS conditions and the enriched library of selected cell-SELEX rounds were prepared for Illumina sequencing as described in Zumrut et al. Briefly, a two-step PCR approach was employed. In the first step, PCR amplicon, specific overhang primers made from Illumina’s overhang adapters, and the SELEX primer (italicized) sequences were used to amplify the sequences. The overhang primers were purchased from IDT, and the sequences were as shown: Forward Amplicon: 5’-TCGTCGGCAGCGTCAGATGTGTATAAGAGACAGATGAGAGCGTCGGTGTGGTA-3’ 5’-GTCTCGTGGGCTCGGAGATGTGTATAAGAGACAGTACTTCCGCACCCTCCTACA-3’, reverse amplicon. The PCR amplicon mixture for the selected SELEX rounds or LIGs contained 10 μL of a 10 nM DNA template in TRIS-HCL buffer, 5 μL of 1 μM forward or reverse amplicon, 25 μL 2X KAPA HiFi HotStart ReadyMix (KAPA Biosystems, KK2601), and 5 μL of DNase-free water in a final reaction volume of 50μL. The number of PCR cycles for the amplification of each sample was optimized, and the chosen cycle amplicon was purified using Ampure XP beads. Following purification of the PCR amplicon product, a second step of PCR, Index PCR, was performed to introduce unique Illumina indices for multiplexing. Each sample was prepared with two different indexes, index 1 (i7) and index 2 (i5), and 2X KAPA HiFi HotStart ReadyMix to generate uniquely tagged libraries pooled together for sequencing. The resulting PCR product was purified and characterized using agarose gel electrophoresis. All samples were submitted to the genomic and epigenomic core facility at Weill-Cornell Medicine (WCM) for high-throughput Illumina sequencing. Four samples were pooled together to maintain high sequence coverage and sequenced using the Illumina NovaSeq 6000 instrument in paired-end mode with 100 cycles as read length.

### FASTAptamer and Galaxy analysis

Initial analysis of the sequenced data from SELEX and LIGS was carried out with the FASTAptamer Bioinformatic toolkit. The FASTAptamer module was used to count and rank sequences in reads per million (RPM), and the output was used in the FASTAptamer-Enrich module to calculate the fold enrichment ratio of sequences in different LIGS conditions. Fold enrichment ratios were calculated as previously described^32^ with some modifications, and RPM_y_/RPM_x_, RPM_Z_/RPM_Y,_ and RPM_Z_/RPM_x_ were obtained, where x represented sequences occurring in the enriched SELEX library, y represented sequences outcompeted using the isotype control, and z represented sequences outcompeted using the chemical ligand against β_2_AR or FLAG mAbs. FASTAptamer-enriched output data were further analyzed using the Galaxy platform (usegalaxy.org). The Galaxy streamline was similar to that in a previous study. The first applied filter of z/x (Z_Ligand_/X_SELEX round_) ≥2 was determined from distinct populations in a scatter plot of fold enrichment against RPM of the sequences from SELEX round 11 and round 14 libraries. Similarly, a second filter of z/y (Z_Ligand_/Y_control_) ≥5 was applied to the initially filtered z/x data to exclude high off-rate and off-target sequences. To remove sequences resulting from membrane disruption, sequences from the propranolol ligand were used as a positive filter for LIGS conditions with isoproterenol, epinephrine, and FLAG mAbs. Similarly, isoproterenol was used as a positive filter for propranolol. Next, we filtered out sequences with reads ≤ 2 to remove sequences resulting from PCR or sequencing errors. After applying this final filter, the sequences were sorted in descending order with the Galaxy sort tool and converted to FASTA-formatted sequences using the Tabular-to-FASTA tool. The obtained FASTA sequences were concatenated head-to-tail and aligned with ClustalW. Further sequence alignment analyses were done with ClustalX 2.1 to identify sequence families.

### Fluorescence-based binding assay

Twenty candidate hits were selected from twenty different families and tested against W9 and HeLa cells. Fluorescently labeled sequences, or random sequences, were prepared by denaturation at 95 °C for 10 minutes and then folded at 4 °C for 45 minutes. W9 and HeLa cells were washed three times each with 3 mL of wash buffer. The initial binding test involved incubating 1.5 × 10^5^ W9 or HeLa cells with a final concentration of 500 nM of each candidate or control sequence at 4 °C for 45 minutes. After incubation, the cells were washed once with 3 mL of the wash buffer, resuspended in 300 μL of wash buffer, and analyzed by flow cytometry to record 10,000 binding events. The percentage of specific binding was calculated as Percent specific binding = [(median fluorescence of aptamer candidate) – (median fluorescence of random)] / (median fluorescence of random) × 100.

### Fluorescence-based apparent affinity analysis

The apparent affinity of aptamers targeting β_2_AR was determined using concentrations ranging from 5 nM to 250 nM. Aptamers, or random sequences, and W9 cells were prepared as mentioned above. Each concentration was incubated with 1.5 × 10^5^ W9 cells at 4 °C for 45 minutes, and after incubation, cells were washed once with 3 mL of wash buffer and resuspended in 300 μL of wash buffer. Binding events were analyzed with flow cytometry, recording 5000 binding events. *K*_*d*_ values were obtained by plotting the specific median fluorescence against concentration in GraphPad Prism, where specific median fluorescence = median fluorescence of aptamer – median fluorescence of random. The normalized specific median fluorescence was analyzed using nonlinear regression (curve fit) and one-site-specific binding.

### Analysis of aptamer specificity

Different specificity assays were performed to determine the specificity for β_2_AR. First, fluorescently labeled aptamers were assessed for their specific binding to W9 cells overexpressing the receptor and the control HeLa cells. Aptamers, or random sequences, were prepared as mentioned above. Subsequently, 250 nM of each aptamer, or random sequence, were incubated with 7.5 × 10^4^ W9 cells or HeLa cells at 25 °C or 37 °C for 45 minutes. After incubation, cells were washed twice with 2 mL of wash buffer and analyzed with flow cytometry, recording 10,000 binding events. Second, W9 cells were treated with 10 μM isoproterenol for 45 minutes at 37 °C. After treatment, cells were stained with aptamers, or random, and FLAG mAbs, or isotype control. The specific median fluorescence was plotted with GraphPad P and analyzed using the non-parametric *t*-test. All experiments were performed in triplicate.

### Fluorescence-based ligand competition assay

CY3 EV β_2_AR 16 aptamer was purchased from IDT and used for competition assays. 4 × 10^5^ β_2_AR-overexpressing W9 cells in 100 μL of cell suspension buffer were pretreated with 1 μM CY3-labeled aptamer, or 8.2 nM FLAG mAbs, for 30 minutes at 4°C, followed by washing before incubation with 250 nM of FAM-labeled aptamers prepared as previously mentioned for an additional 45 minutes at 4°C. After incubation, the cells were washed once with 3 mL wash buffer and resuspended in 300 ul of the wash buffer. Fluorescence intensity was quantified through flow cytometry by recording 5000 events; median fluorescence was normalized to the signals obtained from the binding of FAM-labeled aptamers to untreated W9 cells (serving as the control) and random. The specific median fluorescence values were calculated and graphically represented using GraphPad Prism and employing one-way ANOVA.

### Flow cytometric analysis of β_2_AR internalization

The washed W9 cells were treated with 250 nM of prepared aptamers or a control for 60 minutes at 37°C. After treatment, the cells were placed on ice for 5 minutes, washed once with 2 mL of wash buffer, and then resuspended in 100 μL of 0.25% trypsin-EDTA in Hanks’ Balanced Salt Solution (HBSS) or wash buffer for 5 minutes at 37°C. Trypsin activity was stopped by adding 100 μL of 10% FBS in DMEM buffer. Cells were washed once with 2 mL of wash buffer and resuspended in 300 μL of wash buffer. Internalization was assessed by recording 5000 events using flow cytometry. The normalized median fluorescence was plotted and analyzed with GraphPad Prism.

### Confocal microscopy analysis

W9 cells were treated with 250 nM of prepared aptamers for 60 minutes at 37°C. After treatment, the cells were placed on ice for 5 minutes, washed once with 2 mL of wash buffer, and then resuspended in 2 mL of 1X PBS containing 1 μL of Hoechst nucleus staining dye. Cells were then incubated for 10 minutes at 37 °C. Following incubation, cells were centrifuged and resuspended in 100 μL of 0.25% trypsin-EDTA in HBSS, or wash buffer, and incubated for 5 minutes at 37 °C. Trypsin activity was stopped by adding 100 μL of 10% FBS in DMEM, and cells were washed with 2 mL of wash buffer. The cell pellet was resuspended in 100 μL of HBSS and transferred to a cell dish with a coverslip.

### Microscale Thermophoresis

Microscale Thermophoresis (MST) was used to quantify the binding interaction between Cy5-labeled aptamer and unlabeled purified β_2_-Adrenergic Receptor (Protein). Cy5-labeled aptamers were purchased from IDT, and β_2_AR was purified in the Robert J. Lefkowitz laboratory at Duke University. Cy5-labeled aptamers were maintained at a constant concentration of 1µM, while the binding partner, purified protein, was titrated in 1:1 serial dilutions from 30µM to 9nM. Purified protein was diluted in DMEM buffer containing 0.01% MNG and 0.001% CHS.

Prior to binding, the aptamers were denatured at 95℃ for 7 minutes, followed by folding on ice for 45 minutes. Equal volumes of aptamer and protein serial dilutions were then mixed and incubated for 20 minutes at 25℃ (room temperature). After incubation, the samples were loaded into the capillaries and measured on the Monolith X instrument (NanoTemper) using 1% MST power for β_2_AR16 and 3% MST power for β_2_AR19. Binding curves were generated by plotting the normalized fluorescence values against the logarithm of protein concentration using XY scatter plots in GraphPad Prism. All measurements were performed in triplicate.

### Statistical analysis

All statistical tests and curve fitting (nonlinear regression) were conducted using GraphPad Prism. Statistical comparisons were made with either the non-parametric *t*-test or one-way ANOVA.

## Notes

### Competing Interest Statement

The authors have declared no competing interest.

