## Supplementary material for "Precision Aptamers Against a Native GPCR through Ligand-Guided Selection": SI file: 08 Supplementary information 2 _3_AG.pdf

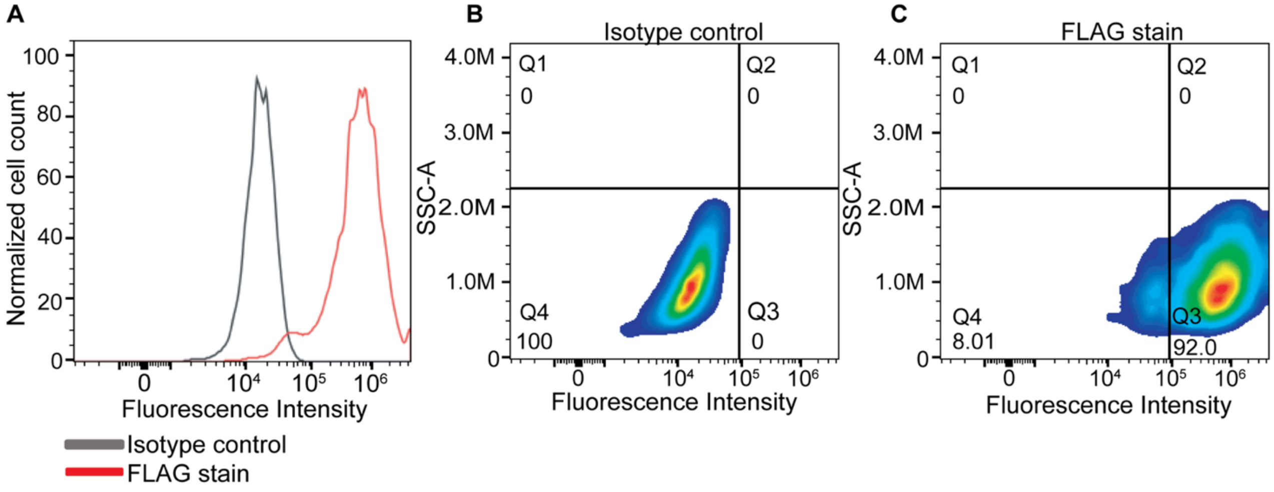


**Figure S1: Monitoring the expression of $\beta_2$AR in W9-overexpressing cells.** A-C) W9 cells were cultured to approximately 90% confluency. The cultured cells were enzymatically detached with trypsin-EDTA in 5 mL of DPBS. Trypsin activity was stopped with 10% DMEM, and the cells were pelleted at 1000 g for 3 minutes. The cells were resuspended in 10 mL of 10% DMEM medium to obtain the cell count. W9 cells in the required concentration were washed 3 times in the SELEX buffer. After washing, the cells were resuspended in cell-suspension buffer to a concentration of 1.0 × $10^6$ cells/95 µL, and 5 µL of the mAbs, or isotype control, were added. The cells were incubated on ice for 30 minutes, washed with 2 ml of the SELEX buffer, and then analyzed using flow cytometry.

**Table S1: Cell-SELEX conditions utilized during the selection process.**

| SELEX rounds | Positive selection plate size | Counter selection plate size | Washing conditions |
|---|---|---|---|
| 1$^{st}$ | 100 mm | - | 9 mL TBS x 1 |
| 2$^{nd}$ | 100 mm | - | 3 mL TBS x 2 |
| 3$^{rd}$ | 100 mm | - | 3 mL TBS x 2 |
| 4$^{th}$ | 100 mm | - | 3 mL TBS x 3 |
| 5$^{th}$ | 100 mm | - | 3 mL TBS X 3 |
| 6$^{th}$ | 100 mm | - | 3 mL TBS X3 |
| 7$^{th}$ | 100 mm | - | 3 mL TBS X3 |
| 8$^{th}$ | 60 mm | - | 3 mL TBS X 3 |
| 9$^{th}$ | 60 mm | - | 3 mL TBS X3 |
| 10$^{th}$ | 60 mm | - | 3 mL TBS X 3 |
| 11$^{th}$ | 60 mm | - | 3 mL TBS X 3 |
| 12$^{th}$ | 60 mm | - | 3 mL TBS X 3 |
| 13$^{th}$ | 60 mm | 60 mm | 3 mL TBS X 3 |
| 14$^{th}$ | 60 mm | 60 mm | 3 mL TBS X 3 |
| 15$^{th}$ | 60 mm | 60 mm | 3 mL TBS X 3 |

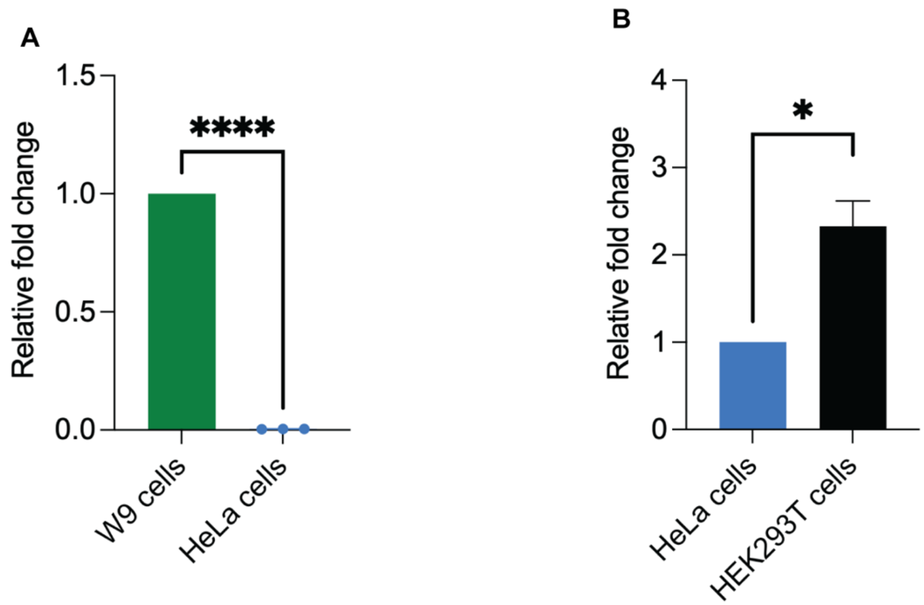


**Figure S2: Quantitative PCR analysis of β2AR expression in different cell lines**. Relative mRNA levels of β2AR were quantified by qPCR in W9, HEK293T, and HeLa cells. Total RNA was extracted, quantified, and reverse transcribed into cDNA. The cDNA was amplified using β2AR-specific primers and detected using the SYBR green detection system. Expression levels were normalized to the housekeeping gene GAPDH and presented as fold-change relative to the control using the □□Ct method. Experiments were conducted in triplicate. Statistical analysis was performed using a nonparametric t-test with significance indicated at $p < 0.05$. A) mRNA expression of β2AR was compared between W9-overexpressing cells and HeLa cells, $p < 0.0001$. B) mRNA expression of β2AR was compared between the endogenously expressing HEK293T cells and HeLa cells endogenously expressing β2AR at very low levels.

**Table S2: FASTAptamer Bioinformatic Count Analysis of Illumina Sequence Data Derived from Libraries Prepared Across Various Rounds of the Cell-SELEX and Ligand-Guided Selection Processes.** Percent enrichment was calculated as (Enrichment × 100). AFL denotes the absence of free ligand (specific sequences). PFL denotes the presence of free ligand (both specific and nonspecific sequences). ISOPRO denotes Isoproterenol, EPI denotes Epinephrine, PROP denotes Propranolol, and FLAG denotes FLAG Monoclonal Antibodies.

| Sample ID | Unique Reads | Total Reads | Complexity (unique/total) | Enrichment (1-complexity) | % Enrichment |
|---|---|---|---|---|---|
| R2 | 28786307 | 32077009 | 0.897 | 0.103 | 10.3 |
| R4 | 16495632 | 18573541 | 0.888 | 0.112 | 11.2 |
| R6 | 18807651 | 21273000 | 0.884 | 0.116 | 11.6 |
| R7 | 31798582 | 38318711 | 0.829 | 0.171 | 17.1 |
| R8 | 1977117 | 3029898 | 0.652 | 0.348 | 34.8 |
| R9 | 20907817 | 71611862 | 0.291 | 0.709 | 70.9 |
| R11 | 1299508 | 48405525 | 0.026 | 0.974 | 97.4 |
| R12 | 370612 | 86194604 | 0.004 | 0.996 | 99.6 |
| R13 | 262269 | 1.01E+08 | 0.002 | 0.998 | 99.8 |
| R14 | 83182 | 51746696 | 0.001 | 0.999 | 99.9 |
| R15 | 30037 | 17769963 | 0.001 | 0.999 | 99.9 |

| | | | | | |
|---|---|---|---|---|---|
| R11_CONTROL_100_AFL | 595015 | 35552935 | 0.016 | 0.984 | 98.4 |
| R11_CONTROL_25_PFL | 1814035 | 66491162 | 0.027 | 0.973 | 97.3 |
| R11_CONTROL_50_AFL | 22116 | 758786 | 0.029 | 0.971 | 97.1 |
| R11_CONTROL_50_PFL | 765249 | 25835912 | 0.029 | 0.971 | 97.1 |
| R11_EPI_100_AFL | 1279474 | 52947646 | 0.024 | 0.976 | 97.6 |
| R11_EPI_25_PFL | 897744 | 25076242 | 0.035 | 0.965 | 96.5 |
| R11_EPI_50_AFL | 264368 | 12987870 | 0.02 | 0.98 | 98 |
| R11_EPI_50_PFL | 792820 | 21588777 | 0.036 | 0.964 | 96.4 |
| R11_FLAG_100_AFL | 929575 | 62663278 | 0.014 | 0.986 | 98.6 |
| R11_FLAG_25_PFL | 1192501 | 43337622 | 0.027 | 0.973 | 97.3 |
| R11_FLAG_50_AFL | 456911 | 26575035 | 0.017 | 0.983 | 98.3 |
| R11_FLAG_50_PFL | 953081 | 37400648 | 0.025 | 0.975 | 97.5 |
| R11_ISOPRO_100_AFL | 1162999 | 48821288 | 0.023 | 0.977 | 97.7 |
| R11_ISOPRO_25_PFL | 909209 | 31183024 | 0.029 | 0.971 | 97.1 |
| R11_ISOPRO_50_AFL | 340306 | 12353823 | 0.027 | 0.973 | 97.3 |
| R11_ISOPRO_50_PFL | 389163 | 12545154 | 0.031 | 0.969 | 96.9 |
| R11_LIB | 1185849 | 33914513 | 0.034 | 0.966 | 96.6 |
| R11_PROP_100_AFL | 667077 | 24169191 | 0.027 | 0.973 | 97.3 |
| R11_PROP_25_PFL | 1753813 | 53830816 | 0.032 | 0.968 | 96.8 |
| R11_PROP_50_AFL | 352558 | 11939058 | 0.029 | 0.971 | 97.1 |
| R11_PROP_50_PFL | 620401 | 17808849 | 0.034 | 0.966 | 96.6 |
| R14 _CONTROL_125_AFL | 215551 | 88921217 | 0.002 | 0.998 | 99.8 |
| R14 _CONTROL_30_PFL | 156388 | 68829661 | 0.002 | 0.998 | 99.8 |
| R14 _CONTROL_60_PFL | 86697 | 32764375 | 0.002 | 0.998 | 99.8 |
| R14_CONTROL_62_5_AFL | 174776 | 72355257 | 0.002 | 0.998 | 99.8 |
| R14 _EPI_125_AFL | 142843 | 60482005 | 0.002 | 0.998 | 99.8 |
| R14 _EPI_30_PFL | 74416 | 29595164 | 0.002 | 0.998 | 99.8 |
| R14 _EPI_60_PFL | 54582 | 20439504 | 0.002 | 0.998 | 99.8 |
| R14 _EPI_62_5_AFL | 38267 | 10399546 | 0.003 | 0.997 | 99.7 |
| R14 _FLAG_125_AFL | 133760 | 44206021 | 0.003 | 0.997 | 99.7 |
| R14 _FLAG_30_PFL | 118072 | 45847483 | 0.002 | 0.998 | 99.8 |
| R14 _FLAG_60_PFL | 142164 | 62296267 | 0.002 | 0.998 | 99.8 |
| R14 _FLAG_62_5_AFL | 130395 | 60812275 | 0.002 | 0.998 | 99.8 |
| R14 _ISOPRO_125_AFL | 118908 | 42319288 | 0.002 | 0.998 | 99.8 |
| R14 _ISOPRO_30_PFL | 84884 | 34225935 | 0.002 | 0.998 | 99.8 |
| R14 _ISOPRO_60_PFL | 71553 | 28784204 | 0.002 | 0.998 | 99.8 |
| R14 _ISOPRO_62_5_AFL | 50119 | 15520146 | 0.003 | 0.997 | 99.8 |
| R14 _LIB | 132718 | 53624765 | 0.002 | 0.998 | 99.8 |
| R14 _PROP_125_AFL | 209461 | 87581535 | 0.002 | 0.998 | 99.8 |
| R14 _PROP_30_PFL | 115894 | 48864890 | 0.002 | 0.998 | 99.8 |
| R14 _PROP_60_PFL | 83116 | 32428005 | 0.002 | 0.998 | 99.8 |
| R14 _PROP_62_5_AFL | 91363 | 32911593 | 0.002 | 0.998 | 99.8 |

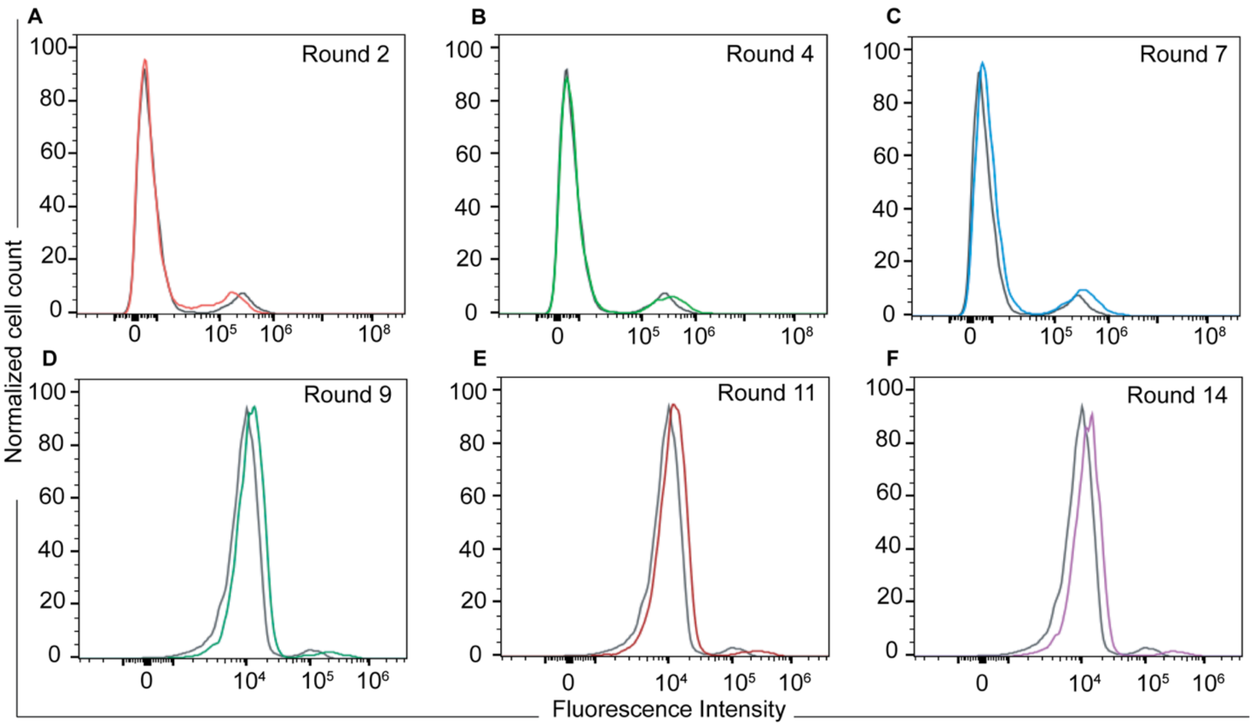


**Figure S2: Progression of the selection**. The progress of SELEX was assessed by incubating 1.5 × $10^5$ W9 cells with 12.5 pmol of the FAM-labeled libraries from various rounds of the cell-SELEX procedure. Changes in median fluorescence intensity were quantified through flow cytometry. No shift in fluorescence intensity was observed in rounds 2-4 (A-B) compared with FAM-labeled round zero library. An increase in fluorescence intensity was observed from round 7 to 14 (C-F).

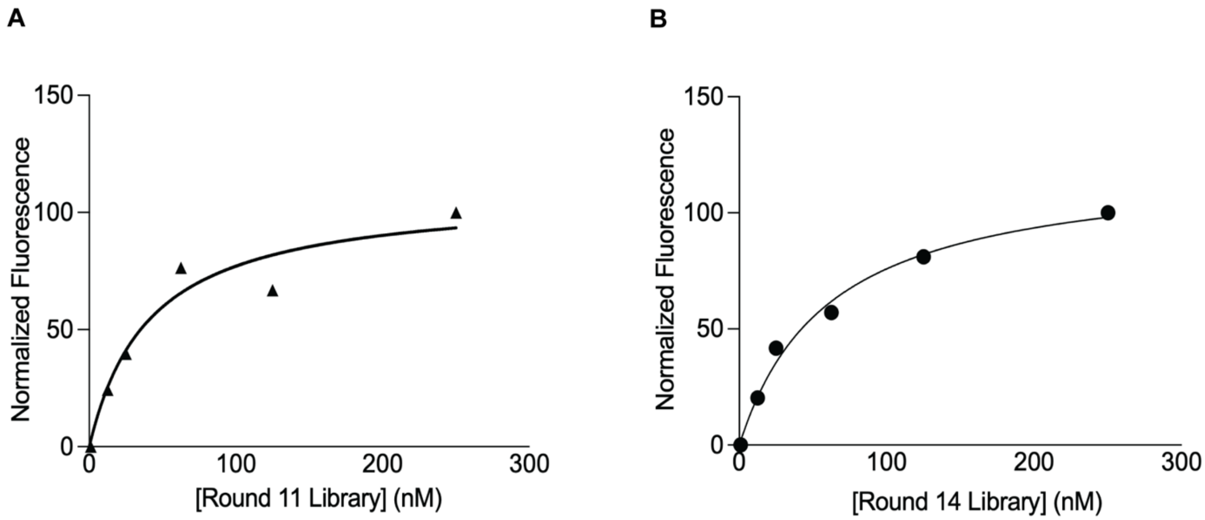


**Figure S3: Apparent affinity determination of the enriched cell-SELEX libraries**. $K_d$ values of enriched SELEX R11 (E) and R14 (F) were determined by incubating 1.5 × $10^5$ W9 cells with concentrations ranging from 1 nM to 250 nM of the respective SELEX libraries at 4 °C for 45 minutes. Library binding was measured by flow cytometry, and the change in fluorescence intensity was plotted and normalized using GraphPad Prism.

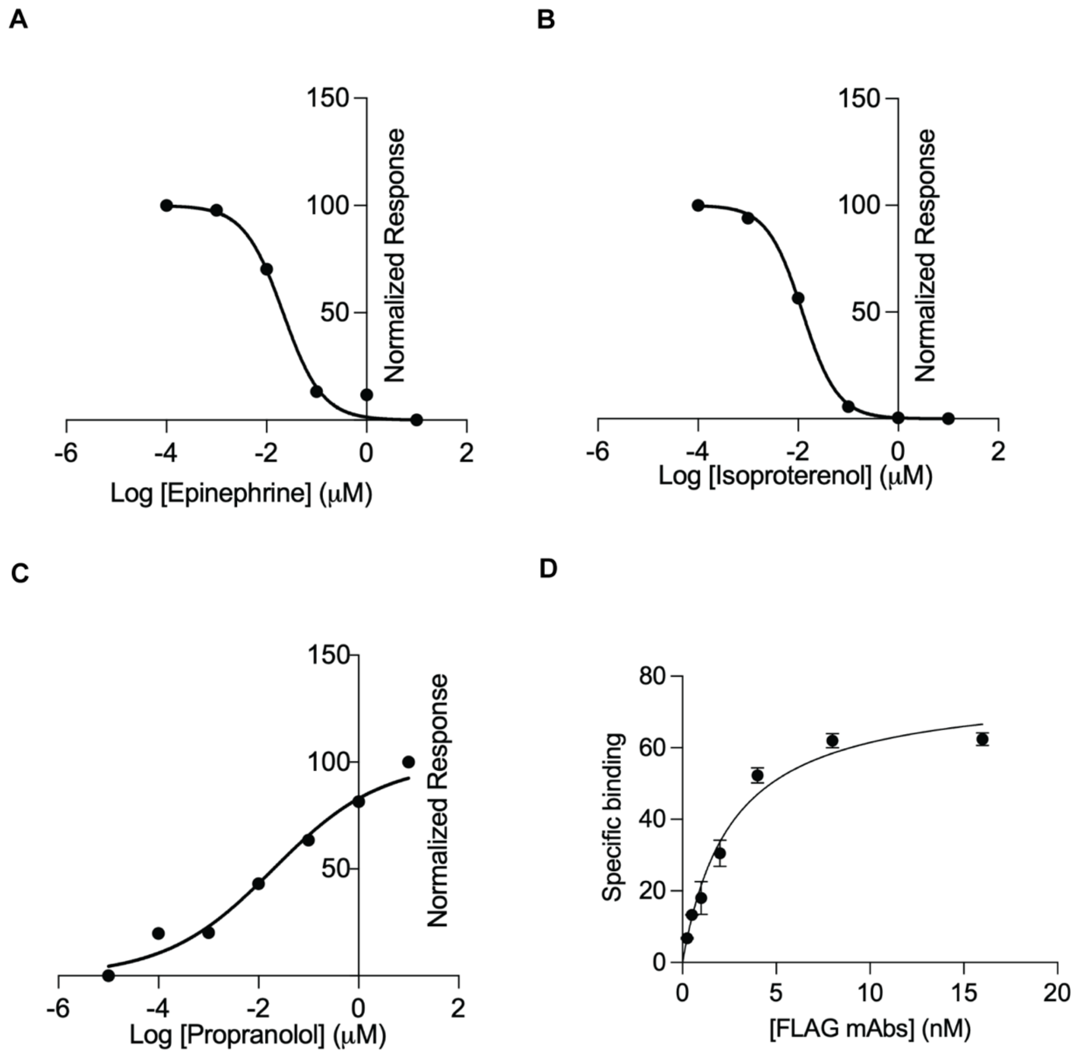


**Figure S4: Characterization of secondary ligands for ligand-guided selection.** A-C) W9 cells were treated with various concentrations (0.1 μM to 10 μM) of epinephrine, isoproterenol, and propranolol, respectively. The effect of ligand concentration on $\beta_2$AR was assessed by staining with FLAG mAbs and plotting the resulting dose-response curve using GraphPad Prism. D) Binding affinity of the FLAG mAbs was determined by incubating W9 cells with concentrations ranging from 0.25 nM to 16 nM of the mAbs at 4°C for 30 minutes. The specific binding for each condition was plotted against the given concentration using GraphPad Prism.

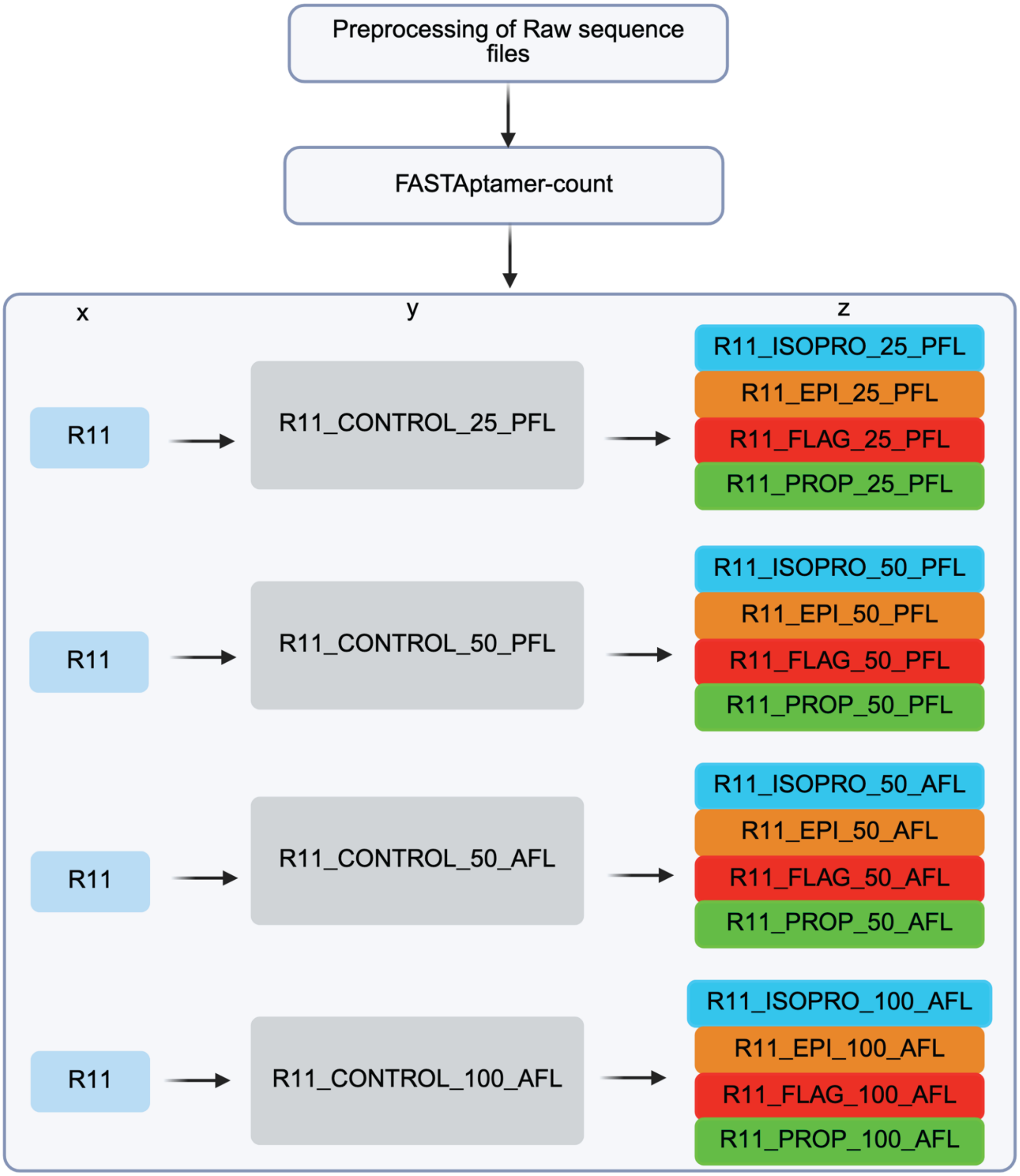


**Figure S5: Bioinformatics analysis of sequenced datasets from ligand-guided selection using R11-enriched SELEX library.** A) Raw data were pre-processed to eliminate primer regions and exclude low-quality reads with a score below 20 at any single position. The pre-processed sequences were then ranked by reads per million (RPM) using the FASTAptamer count module. The output generated by the FASTAptamer module was subsequently employed to calculate the fold enrichment ratios for sequences in the enriched SELEX round 11 (R11), the isotype control ligand (denoted as y), and the specific ligands (ISOPRO, EPI, PROP, and FLAG), denoted as z. The workflow of this analysis is depicted in the accompanying scheme. ISOPRO denotes Isoproterenol; EPI denotes Epinephrine; PROP denotes Propranolol; and FLAG denotes FLAG monoclonal antibodies.

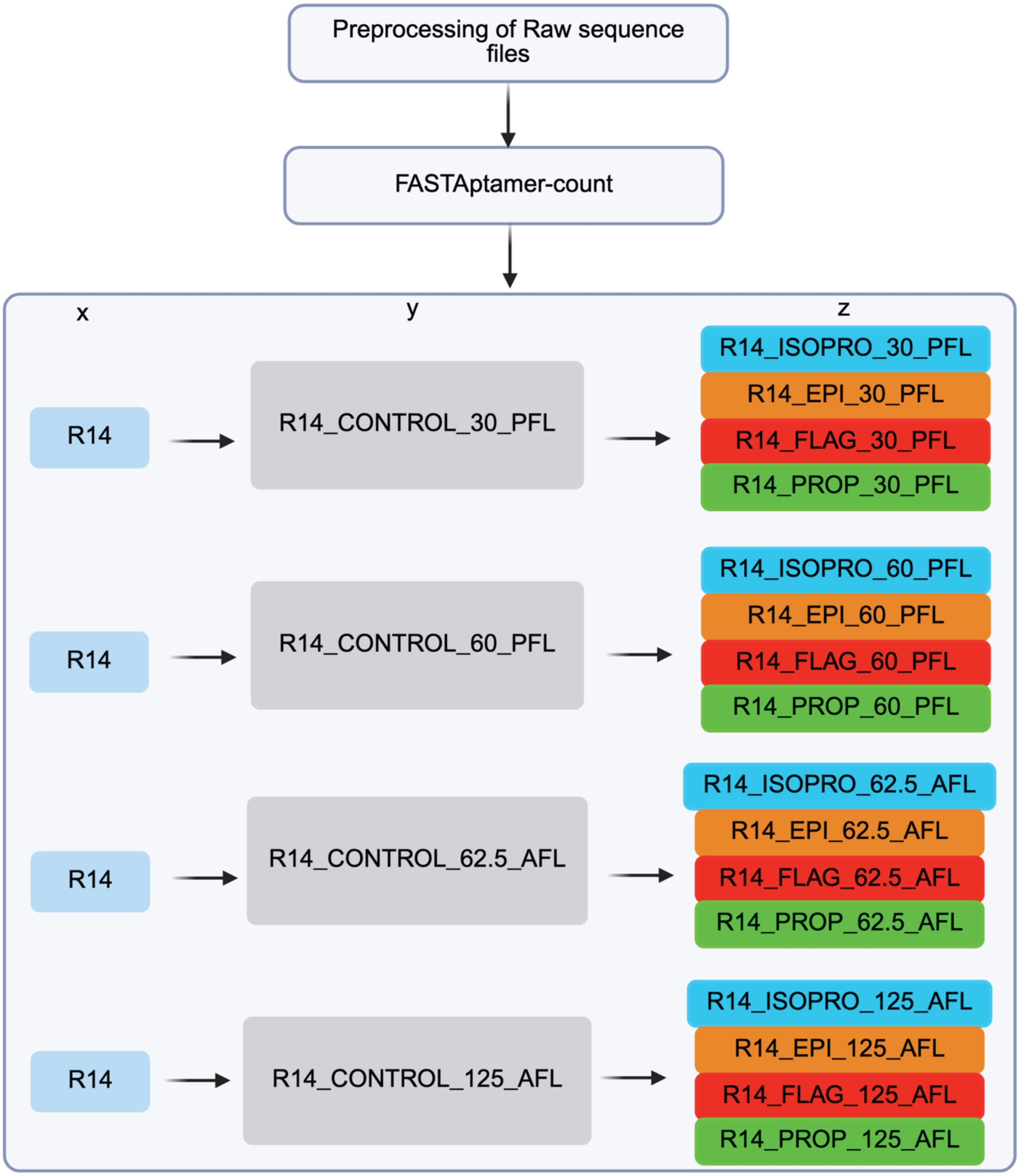


**Figure S6: Bioinformatics analysis of sequenced datasets from ligand-guided selection using R14-enriched SELEX library.** A) Raw data were preprocessed to eliminate primer regions and exclude low-quality reads with a score below 20 at any single position. The preprocessed sequences were then ranked by reads per million (RPM) using the FASTAptamer count module. Output generated by the FASTAptamer module was subsequently employed to calculate the fold enrichment ratios for sequences in the enriched SELEX round 14 (x), the isotype control ligand (denoted as y), and the specific ligands (ISOPRO, EPI, PROP, and FLAG) denoted as z. The workflow of this analysis is depicted in the accompanying scheme. ISOPRO denotes Isoproterenol; EPI denotes Epinephrine; PROP denotes Propranolol; and FLAG denotes FLAG monoclonal antibodies.

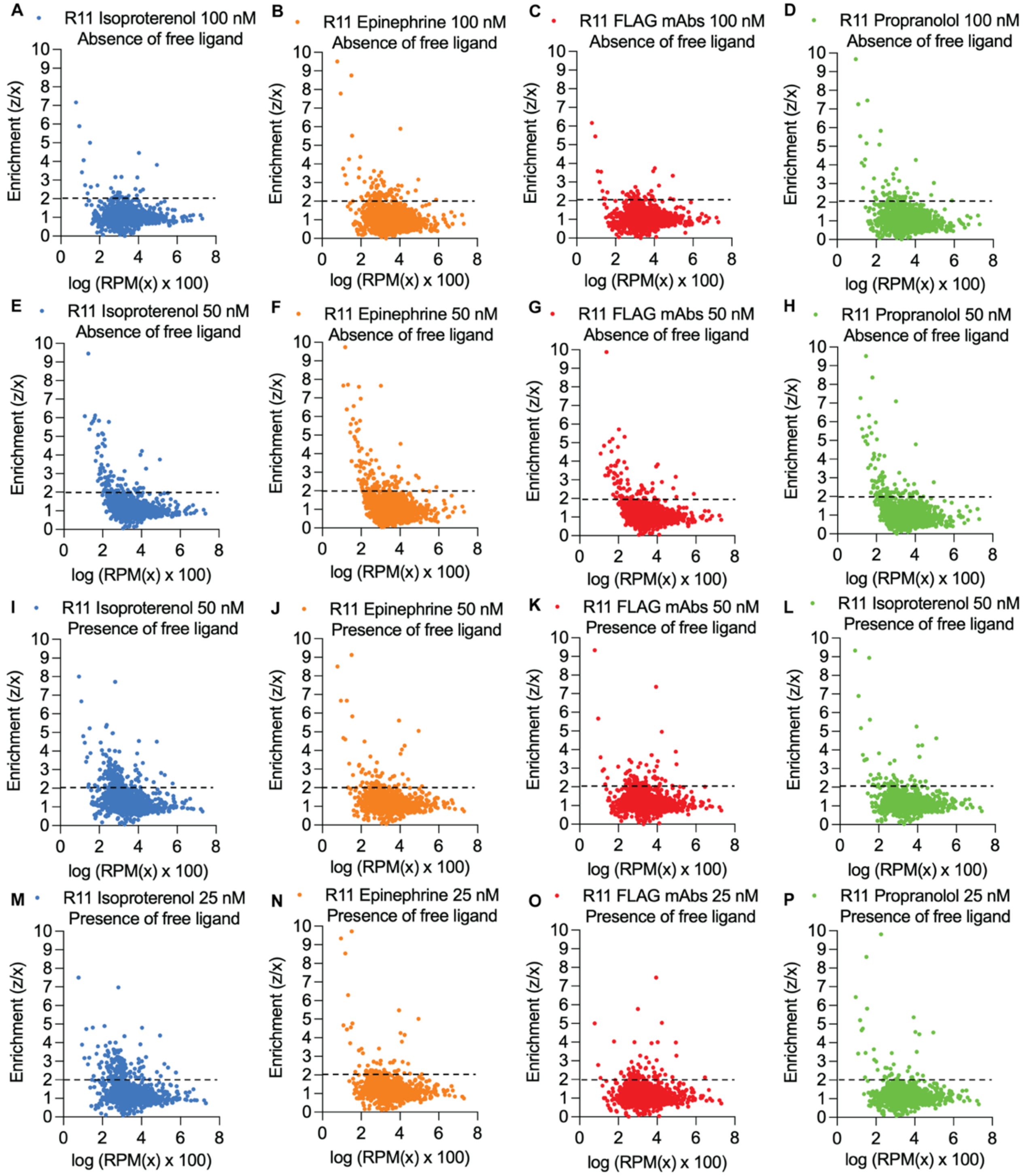


**Figure S7: Scatter plot of the enrichment analysis for each specific ligand condition based on the abundance of sequences in the enriched SELEX library (R11).** The z/x cutoff value was set at fold enrichment ≥2 (black dashes) based on the distribution of sequences in the plot. Sequences with lower fold enrichment (below the line) were discarded to identify sequences specific to $\beta_2$AR. The absence of free ligand indicates bound sequences, while the presence of free ligand shows bound sequences, unbound sequences, and off-target sequences.

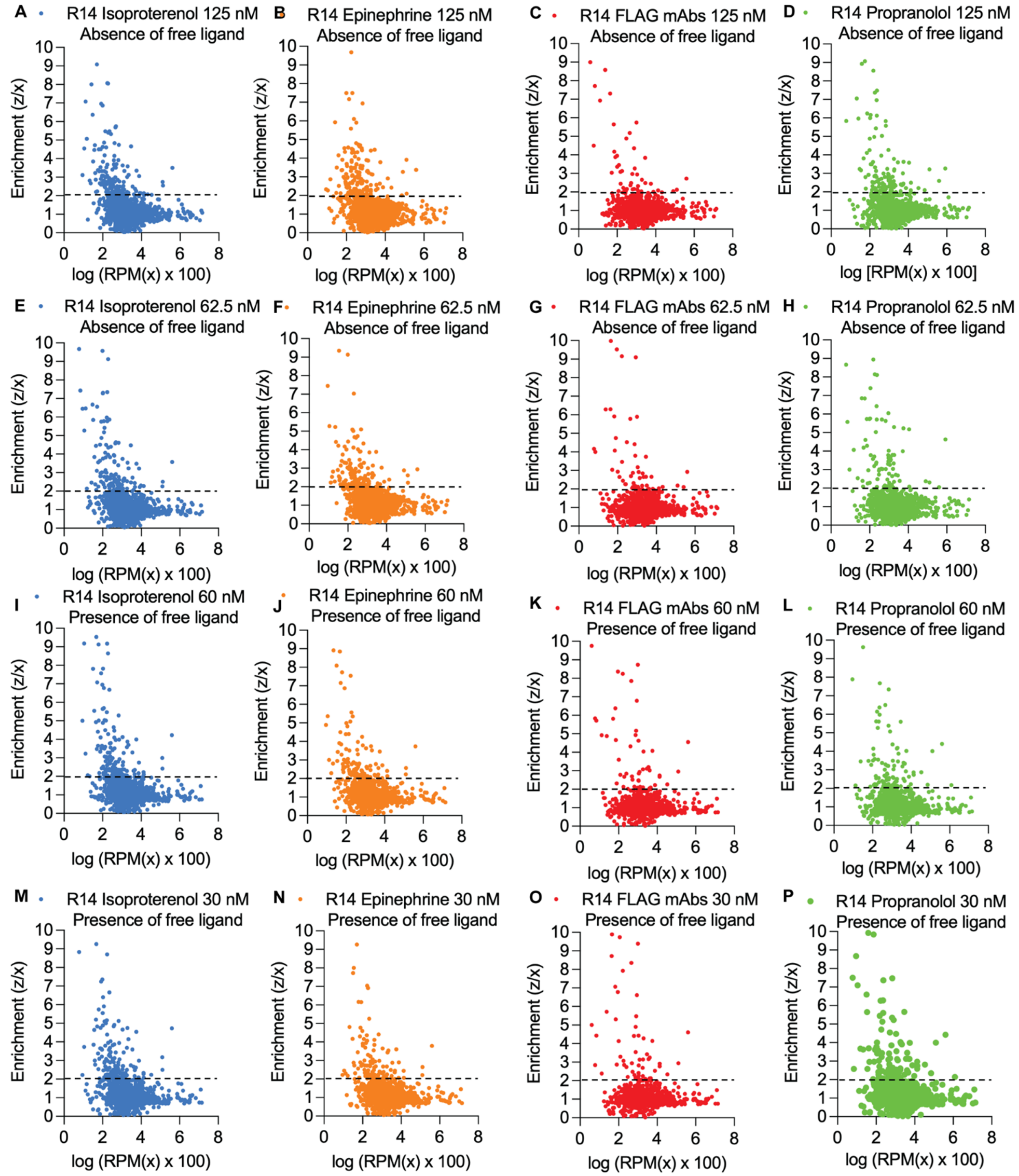


**Figure S8: Scatter plot of enrichment analysis for each specific ligand condition based on the abundance of sequences in the enriched SELEX library (R14).** The z/x cutoff value was set at fold enrichment ≥2 (black dashes) based on the distribution of sequences in the plot. Sequences with lower fold enrichment (below the line) were discarded to identify sequences specific to β$_2$AR. The absence of free ligand indicates bound sequences, while the presence of free ligand shows bound sequences, unbound sequences, and off-target sequences.

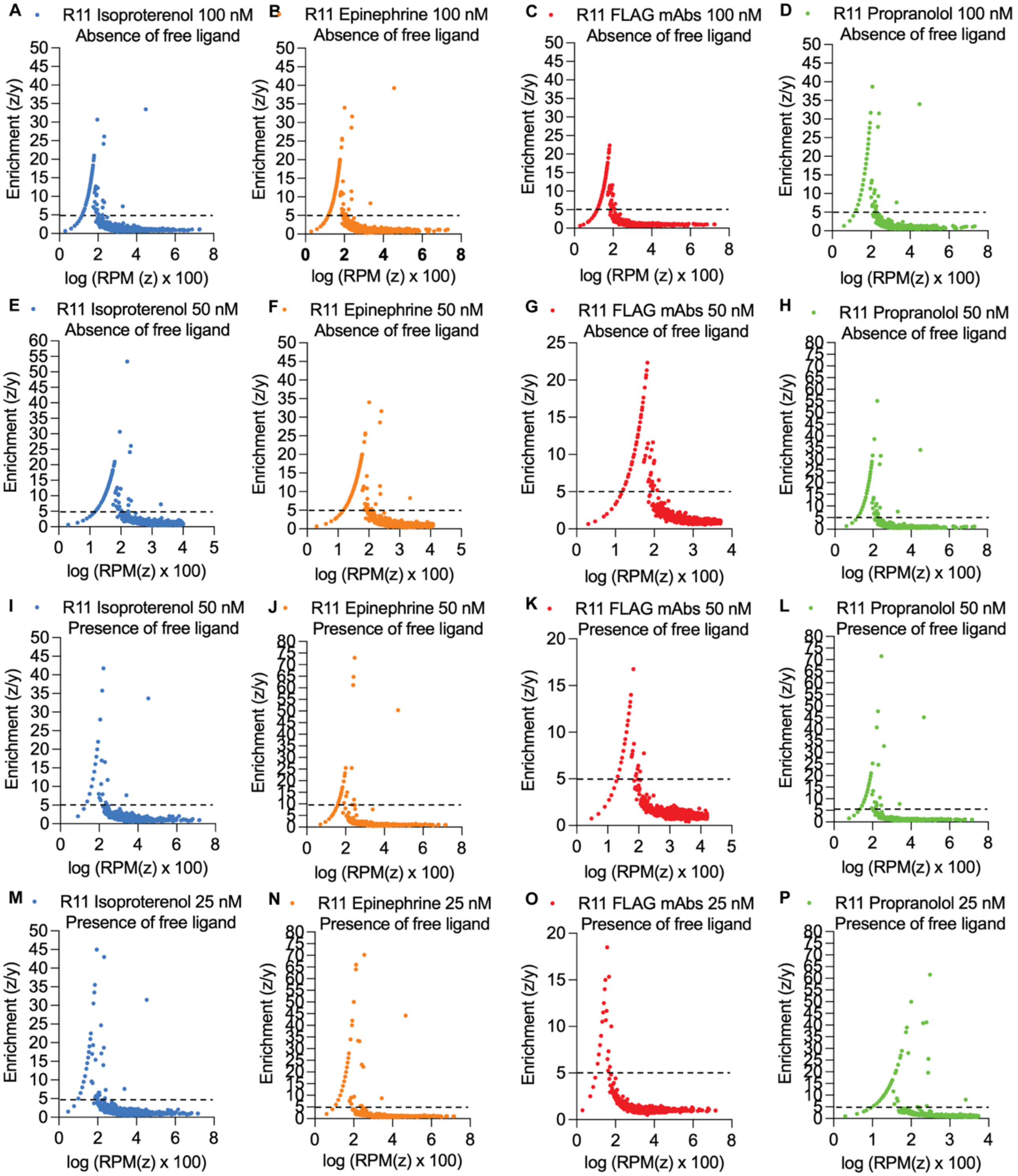


**Figure S9: The enrichment analysis for each specific ligand condition based on the abundance of sequences in the different LIGS sequenced pools (R11).** The z/y cutoff value was set at fold enrichment ≥5 (black dashes) based on the distribution of sequences in the plot. Sequences with lower fold enrichment (below the line) were discarded to identify sequences specific to $\beta_2$AR. The absence of free ligand indicates bound sequences, while the presence of free ligand shows bound sequences, unbound sequences, and off-target sequences.

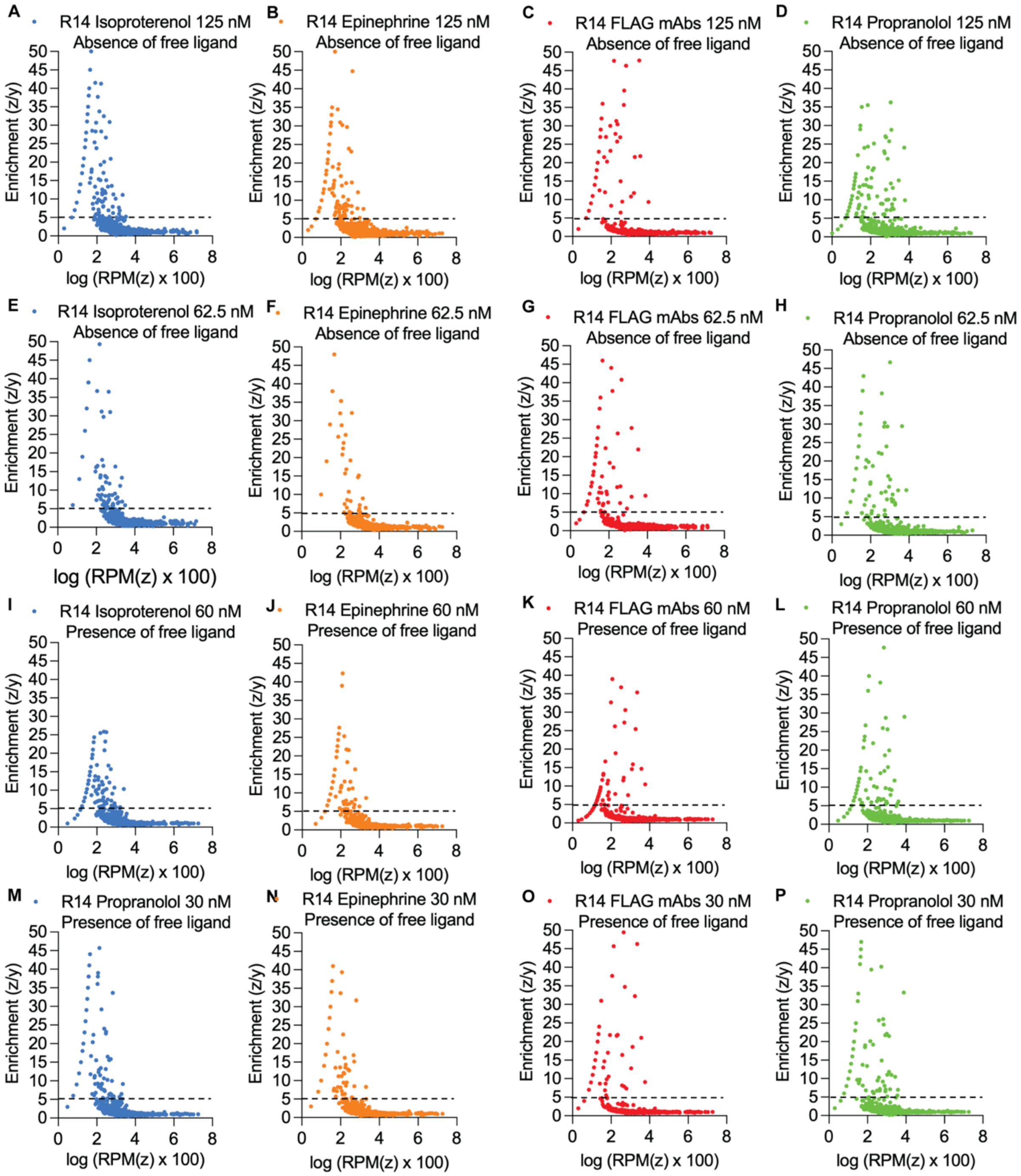


**Figure S10: Scatter plot of the enrichment analysis for each specific ligand condition based on the abundance of sequences in the different LIGS sequenced pools (R14).** The z/y cutoff value was set at fold enrichment ≥5 (black dashes) based on the distribution of sequences in the plot. Sequences with lower fold enrichment (below the line) were discarded to identify sequences specific to β$_2$AR. The absence of free ligand indicates bound sequences, while the presence of free ligand shows bound sequences, unbound sequences, and off-target sequences.

**Table S3: Summary of the number of sequence lines retained after filtering and comparison within the FASTAptamer-enriched data, as analyzed on the GALAXY bioinformatics platform.** AFL denotes the absence of free ligand (specific sequences). PFL denotes the presence of free ligand (both specific and nonspecific sequences). ISOPRO denotes Isoproterenol, EPI denotes Epinephrine, PROP denotes Propranolol, and FLAG denotes FLAG monoclonal antibodies.

| Sample No | File Name | Total no of sequences | No of sequences after filtering z/x ≥2 | No of sequences after filtering z/y ≥5 |
|---|---|---|---|---|
| 1 | R11_vs_R11_CONTROL_25_PFL_vs_R11_ISOPRO_25_PFL | 1,900,000 | 16,234 | 1,031 |
| 2 | R11_vs_R11_CONTROL_50_PFL_vs_R11_ISOPRO_50_PFL | 1,100,000 | 19,987 | 880 |
| 3 | R11_vs_R11_CONTROL_50_AFL_vs_R11_ISOPRO_50_AFL | 961,000 | 16,765 | 12 |
| 4 | R11_vs_R11_CONTROL_100_AFL_vs_R11_ISOPRO_100_AFL | 1,400,000 | 13,293 | 354 |
| 5 | R14_vs_R14_CONTROL_30_PFL_vs_R14_ISOPRO_30_PFL | 179,300 | 7,617 | 1,474 |
| 6 | R14_vs_R14_CONTROL_60_PFL_vs_R14_ISOPRO_60_PFL | 120,900 | 6,422 | 444 |
| 7 | R14_vs_R14_CONTROL_62_5_AFL_vs_R14_ISOPRO_62_5_AFL | 162,900 | 8,508 | 1,924 |
| 8 | R14_vs_R14_CONTROL_125_AFL_vs_R14_ISOPRO_125_AFL | 229,800 | 7,955 | 1,571 |
| 9 | R11_vs_R11_CONTROL_25_PFL_vs_R11_EPI_25_PFL | 1,500,000 | 15,957 | 1,112 |
| 10 | R11_vs_R11_CONTROL_50_PFL_vs_R11_EPI_50_PFL | 932,000 | 13,647 | 440 |
| 11 | R11_vs_R11_CONTROL_50_AFL_vs_R11_EPI_50_AFL | 1,300,000 | 12,344 | 15 |
| 12 | R11_vs_R11_CONTROL_100_AFL_vs_R11_EPI_100_AFL | 1,500,000 | 18,118 | 478 |
| 13 | R14_vs_R14_CONTROL_30_PFL_vs_R14_EPI_30_PFL | 169,500 | 7,033 | 1,341 |
| 14 | R14_vs_R14_CONTROL_60_PFL_vs_R14_EPI_60_PFL | 118,000 | 9,984 | 907 |
| 15 | R14_vs_R14_CONTROL_62_5_AFL_vs_R14_EPI_62_5_AFL | 153,200 | 7,895 | 1,890 |
| 16 | R14_vs_R14_CONTROL_125_AFL_vs_R14_EPI_125_AFL | 241,700 | 6,255 | 1,135 |
| 17 | R11_vs_CONTROL_25_PFL_vs_R11_FLAG_25_PFL | 2,100,00 | 11,316 | 429 |
| 18 | R11_vs_R11_CONTROL_50_PFL_vs_R11_FLAG_50_PFL | 1,400,000 | 9,959 | 240 |
| 19 | R11_vs_R11_CONTROL_50_AFL_vs_R11_FLAG_50_AFL | 1,000,000 | 13,034 | 7 |
| 20 | R11_vs_R11_CONTROL_100_AFL_vs_R11_FLAG_100_AFL | 1,300,000 | 12,025 | 398 |

| | | | | |
|---|---|---|---|---|
| 21 | R14_vs_R14_CONTROL_30_PFL_vs_R14_FLAG_30_PFL | 197,000 | 6,082 | 461 |
| 22 | R14_vs_R14_CONTROL_60_PFL_vs_R14_FLAG_60_PFL | 155,100 | 4,710 | 76 |
| 23 | R14_vs_R14_CONTROL_62_5_AFL_vs_R14_FLAG_62_5_AFL | 210,800 | 4,962 | 764 |
| 24 | R14_vs_R14_CONTROL_125_AFL_vs_R14_FLAG_125_AFL | 238,100 | 5,607 | 960 |
| 25 | R11_vs_R11_CONTROL_25_PFL_vs_R11_PROP_25_PFL | 2,400,000 | 13,762 | 424 |
| 26 | R11_vs_R11_CONTROL_50_PFL_vs_R11_PROP_50_PFL | 1,200,000 | 23,881 | 547 |
| 27 | R11_vs_R11_CONTROL_50_AFL_vs_R11_PROP_50_AFL | 986,900 | 13,579 | 10 |
| 28 | R11_vs_R11_CONTROL_100_AFL_vs_R11_PROP_100_AFL | 1,100,000 | 15,436 | 783 |
| 29 | R14_vs_R14_CONTROL_30_PFL_vs_R14_PROP_30_PFL | 197,100 | 6,855 | 519 |
| 30 | R14_vs_R14_CONTROL_60_PFL_vs_R14_PROP_60_PFL | 130,500 | 5,448 | 240 |
| 31 | R14_vs_R14_CONTROL_62_5_AFL_vs_R14_PROP_62_5_AFL | 190,900 | 4,852 | 822 |
| 32 | R14_vs_R14_CONTROL_125_AFL_vs_R14_PROP_125_AFL | 276,100 | 4,457 | 573 |

**Table S4: Selected hit candidates for initial screening.** Twenty hit sequences were selected from 20 prominent sequence families. In the ligand condition of occurrence, ISOPRO denotes Isoproterenol, EPI denotes Epinephrine, PROP denotes Propranolol, and FLAG denotes FLAG monoclonal antibodies.

| Candidate | Sequence variable region | Ligand condition of occurrence | Enrichment (z/y) | Length |
|---|---|---|---|---|
| EV $\beta_2$AR 1 | CAAGTAGGACTCAGCAGGTTCGGCACGATCCATAAAAA | FLAG | 293 | 38 |
| EV $\beta_2$AR 2 | ATCAGCCCCTGCCCGGAAGTCAAGCTTGCACCTTTGA | Epi | 48 | 37 |
| EV $\beta_2$AR 3 | TCGGCACGGTTTTGTTCATCTACTCGATATACTCTA | Isopro | 130 | 36 |
| EV $\beta_2$AR 4 | ACCCGATGTAGTACAGCTAAGAACCGCGCTTCATCTC | Isopro | 58 | 37 |
| EV $\beta_2$AR 5 | AAGCGTTGGCGTCTTTCGAGTTCCTCCGGTAGCACCG | Isopro | 19 | 37 |
| EV $\beta_2$AR 6 | CACCGCATTTGTTCTGCTTAATAGCTCTCCACGGTAAA | Isopro | 58 | 38 |
| EV $\beta_2$AR 7 | CACCTCTTCTGTTCTGCTTGAAAGCTCTCTCTGGG | Epi | 38 | 35 |
| EV $\beta_2$AR 8 | ATGCTATCTGGTTACGCTTTCACACTACACCGCCGAC | Epi | 48 | 37 |
| EV $\beta_2$AR 9 | GACTGCAGAGACTGCTCGCCGAAACGCGTCGTAACTC | Epi | 23 | 37 |
| EV $\beta_2$AR 10 | GCCGATGGCGGCGTAATCTACGGTTGAATAAATTCGG | Epi | 29 | 37 |
| EV $\beta_2$AR 11 | CCACATGGCCGGGCGTAATTCTCGTCGCCTATCACT | Isopro | 26 | 36 |
| EV $\beta_2$AR 12 | TTCTTGCCACTGCCTTGGACCACCTAGCGGTAATCGC | Epi | 29 | 37 |

| | | | | |
|---|---|---|---|---|
| EV $\beta_2$AR 13 | AACATCGATCGGAAGCCGCCAAGGTTATCAGCCGATT | Prop | 73 | 37 |
| EV $\beta_2$AR 14 | TCGGTCCCCCCGTACCCATCGCATCTCTGATAGGTC | Prop | 50 | 36 |
| EV $\beta_2$AR 15 | GCCGGGACTCTGTATCCCCGACGGCACATGTGCGTAT | Isopro | 19 | 37 |
| EV $\beta_2$AR 16 | ACTCAGCTGGCTAAGTTATTCGCCCAAGTTCCGCCAC | Isopro and Epi | 39 | 37 |
| EV $\beta_2$AR 17 | TGGCGACCCAATCCGGCTAGTAAACGTAGTCCATGTT | FLAG mAbs | 10 | 37 |
| EV $\beta_2$AR 18 | ACCCGATGTAGTACAGCTAAGTACCGTGTTTCATCTC | Epi | 12.6 | 37 |
| EV $\beta_2$AR 19 | GAACCCGCCAGGCCGAATGTCAGTCGTGCACTTTTTC | Isopro | 13 | 37 |
| EV $\beta_2$AR 20 | GCATCGATCGGAAGTCTCCAAGGTTATCAGCCCATT | Isopro and Epi | 21.5 | 36 |

**A**

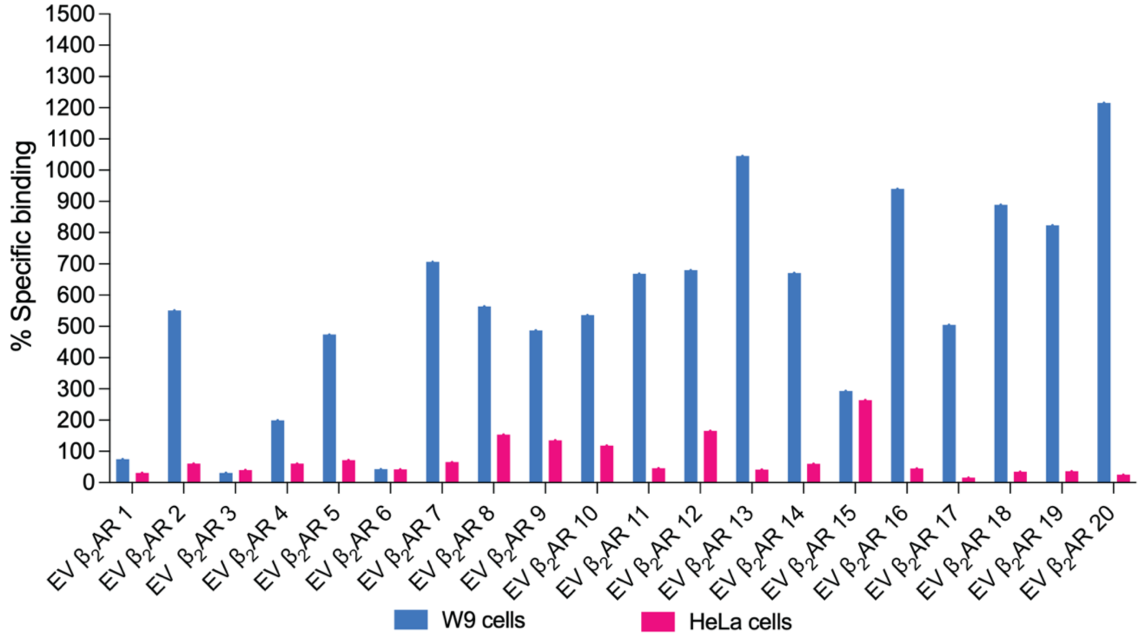


**B**

| Name | Hit candidates | $K_d$ value (nM) |
| --- | --- | --- |
| EV $\beta_2$AR 10 | ---GCCGATGGC--GGCGTAATCTACGGTTGAATAAATTCGG---- | 148.0 |
| EV $\beta_2$AR 11 | ---CCACATGGCCGGGCGTAATTCTCG-TCGCCTATCACT------ | 342.3 |
| EV $\beta_2$AR 2 | ---ATCAGCCCCTGCCCGGAAGTCAAGCTTGCACCTTTGA------ | 335.1 |
| EV $\beta_2$AR 19 | ---GAACCCGCCAGGCCGAATGTCAGTCGTGCACTTTTTC------ | 84.0 |
| EV $\beta_2$AR 13 | AACATCGATCGGAAGCCGCCAAGGTTATCAGCCGATT--------- | 780.4 |
| EV $\beta_2$AR 20 | -GCATCGATCGGAAGTCTCCAAGGTTATCAGCCCATT--------- | 112.2 |
| EV $\beta_2$AR 16 | ---------ACTCAGCTGGCTAAGTTATTCGCCCAAGTTCCGCCAC | 119.4 |
| EV $\beta_2$AR 6 | --CACCGCATTTGTTCTGCTTAATAGCTCTCCACGGTAAA------ | ND |
| EV $\beta_2$AR 7 | --CACCTCTTCTGTTCTGCTTGAAAGCTCTCTCTGGG--------- | 388.7 |
| EV $\beta_2$AR 4 | --ACCCGATGTAGTACAGCTAAGAACCGCGCTTCATCTC------- | ND |
| EV $\beta_2$AR 18 | --ACCCGATGTAGTACAGCTAAGTACCGTGTTTCATCTC------- | 129.5 |
| EV $\beta_2$AR 17 | --TGGCGACCCAATCCGGCTAGTAAACGTAGTCCATGTT------- | 275.5 |
| EV $\beta_2$AR 15 | ------GCCGGGACTCTGTATCCC-CGACGGCACATGTGCGTAT-- | ND |
| EV $\beta_2$AR 1 | ----CAAGTAGGACTCAGCAGGTTCGGCACGATCCATAAAAA---- | ND |
| EV $\beta_2$AR 3 | ----TCGGCACGGTTTTGTTCATCTACTCGATATACTCTA------ | ND |
| EV $\beta_2$AR 5 | --AAGCGTTGGCGTCTTTCGAGTTCCTCCGGTAGCACCG------- | 202.0 |
| EV $\beta_2$AR 12 | -TTCTTGCCACTGCCTTGGACCACCTAGCGGTAATCGC-------- | 123.1 |
| EV $\beta_2$AR 8 | ----ATGCTATCTGGTTACGCTTTCACACTACACCGCCGAC----- | 201.4 |
| EV $\beta_2$AR 14 | ----TCGGTCCCCCCGTACCCATCGCATCTCTGATAGGTC------ | 292.4 |
| EV $\beta_2$AR 9 | GACTGCAGAGACTGCTCGCCGAAACGCGTCGTAACTC--------- | 609.9 |

**Figure S12: Binding affinity assessment of β2AR hit candidates.** A) Initial binding analysis was conducted using a final concentration of 500 nM for each fluorescently labeled hit aptamer candidate or random, which was incubated with W9 or HeLa cells for 45 minutes at 4°C. Subsequently, binding was evaluated by flow cytometry with 10,000 events recorded. The percent specific binding of each candidate was calculated as percent specific binding = [(median fluorescence of aptamer candidate –

median fluorescence of random) / (median fluorescence of random)] × 100. Data were plotted using GraphPad Prism via XY analysis. B) A series of concentrations ranging from 5 nM to 250 nM of the fluorescently labeled hit aptamer candidate or random was incubated with 1.5 × 105 W9 cells for 45 minutes at 4°C. Differential binding was quantified by flow cytometry, recording 5000 events to determine the normalized median fluorescence (Normalized median fluorescence = median fluorescence of aptamer candidate□□ median fluorescence of random), which was subsequently analyzed with GraphPad Prism using a nonlinear, one-site specific binding model. ND equals not determined.

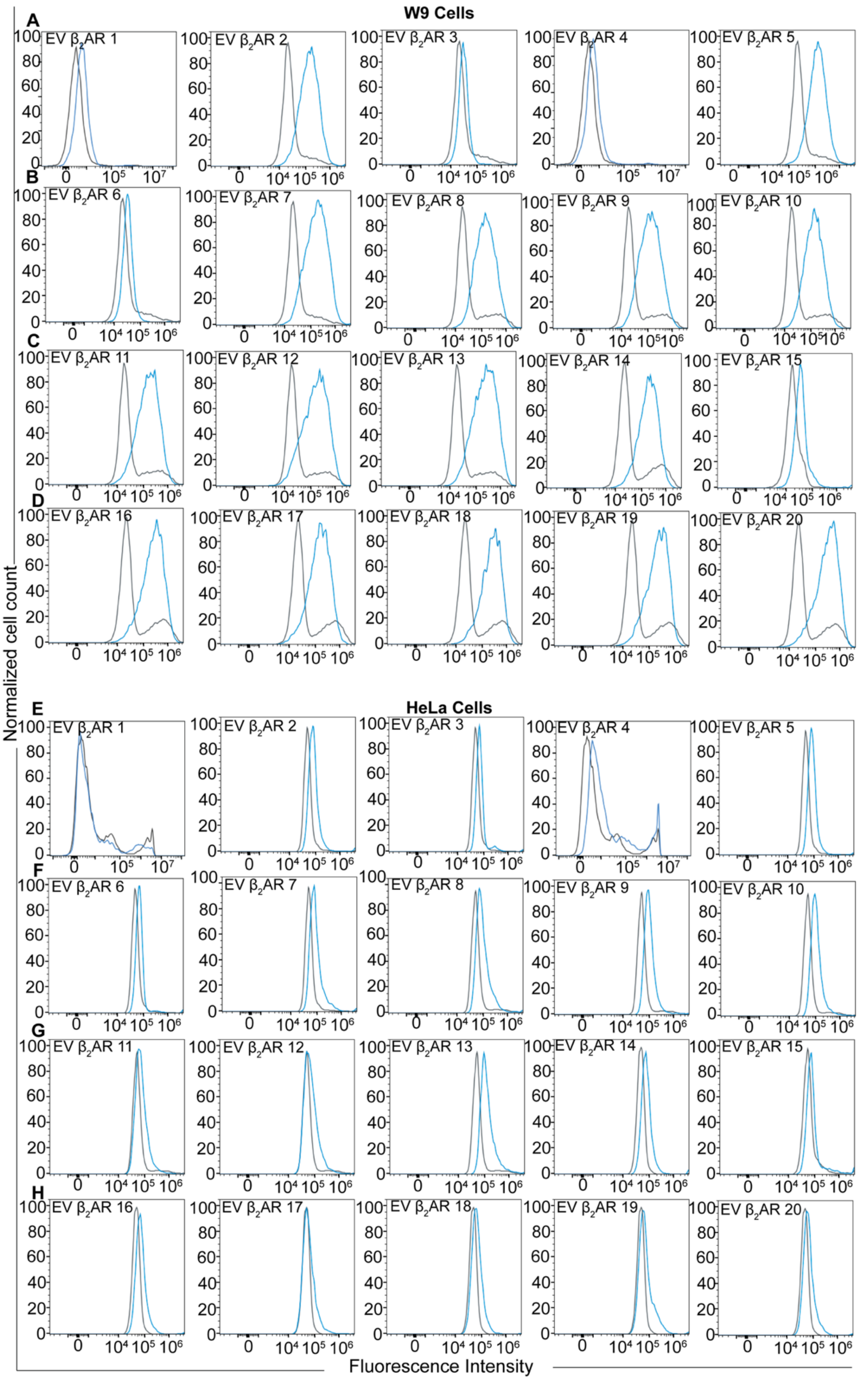
W9 Cells
HeLa Cells
A
B
C
D
E
F
G
H
EV β2AR 1
EV β2AR 2
EV β2AR 3
EV β2AR 4
EV β2AR 5
EV β2AR 6
EV β2AR 7
EV β2AR 8
EV β2AR 9
EV β2AR 10
EV β2AR 11
EV β2AR 12
EV β2AR 13
EV β2AR 14
EV β2AR 15
EV β2AR 16
EV β2AR 17
EV β2AR 18
EV β2AR 19
EV β2AR 20
Normalized cell count
Fluorescence Intensity

**Figure S11: Whole-cell fluorescence-based screening of aptamers against $\beta_2$AR using W9 cells and HeLa cells.** The binding of each FAM-labeled hit candidate was evaluated using W9 and HeLa cells. A total of 1.5 x $10^5$ cells were incubated with 500 nM of each candidate at 4 °C for a duration of 45 minutes. Subsequently, the cells were washed with 2 mL of wash buffer and resuspended in 300 µL of the same buffer. Flow cytometry was employed to record and analyze 10,000 events per sample. Flow cytometry data analysis generated histograms in which the Y-axis represents the normalized cell count versus the change in fluorescence intensity (x-axis). The gray histogram indicates the binding of the negative control (random), while the blue histograms represent the binding of various hit candidates.

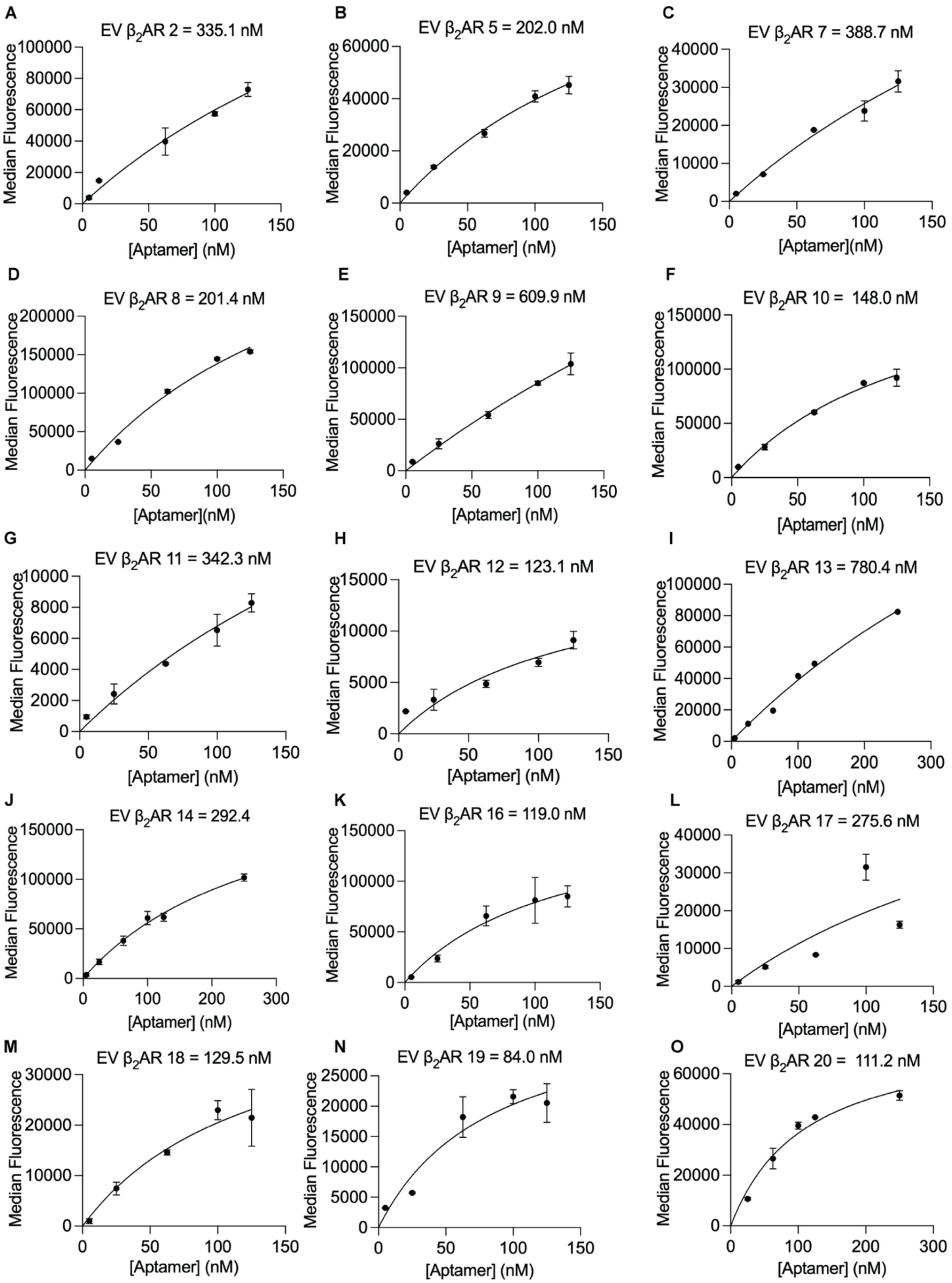
A
EV β2AR 2 = 335.1 nM
B
EV β2AR 5 = 202.0 nM
C
EV β2AR 7 = 388.7 nM
D
EV β2AR 8 = 201.4 nM
E
EV β2AR 9 = 609.9 nM
F
EV β2AR 10 = 148.0 nM
G
EV β2AR 11 = 342.3 nM
H
EV β2AR 12 = 123.1 nM
I
EV β2AR 13 = 780.4 nM
J
EV β2AR 14 = 292.4
K
EV β2AR 16 = 119.0 nM
L
EV β2AR 17 = 275.6 nM
M
EV β2AR 18 = 129.5 nM
N
EV β2AR 19 = 84.0 nM
O
EV β2AR 20 = 111.2 nM
Median Fluorescence
[Aptamer] (nM)

**Figure S12: Determining the apparent affinity of $\beta_2$AR hit candidates.** W9 cells were incubated with the fluorescently labeled hit aptamer candidate at concentrations ranging from 5 nM to 250 nM, or with a random sequence, for 45 minutes at 4°C. Differential binding was quantitatively assessed by flow cytometry, recording 5000 events to determine the specific median fluorescence (the median fluorescence of the aptamer candidate minus that of the random sequence). Data were subsequently analyzed using GraphPad Prism employing a nonlinear, one-site specific binding model.

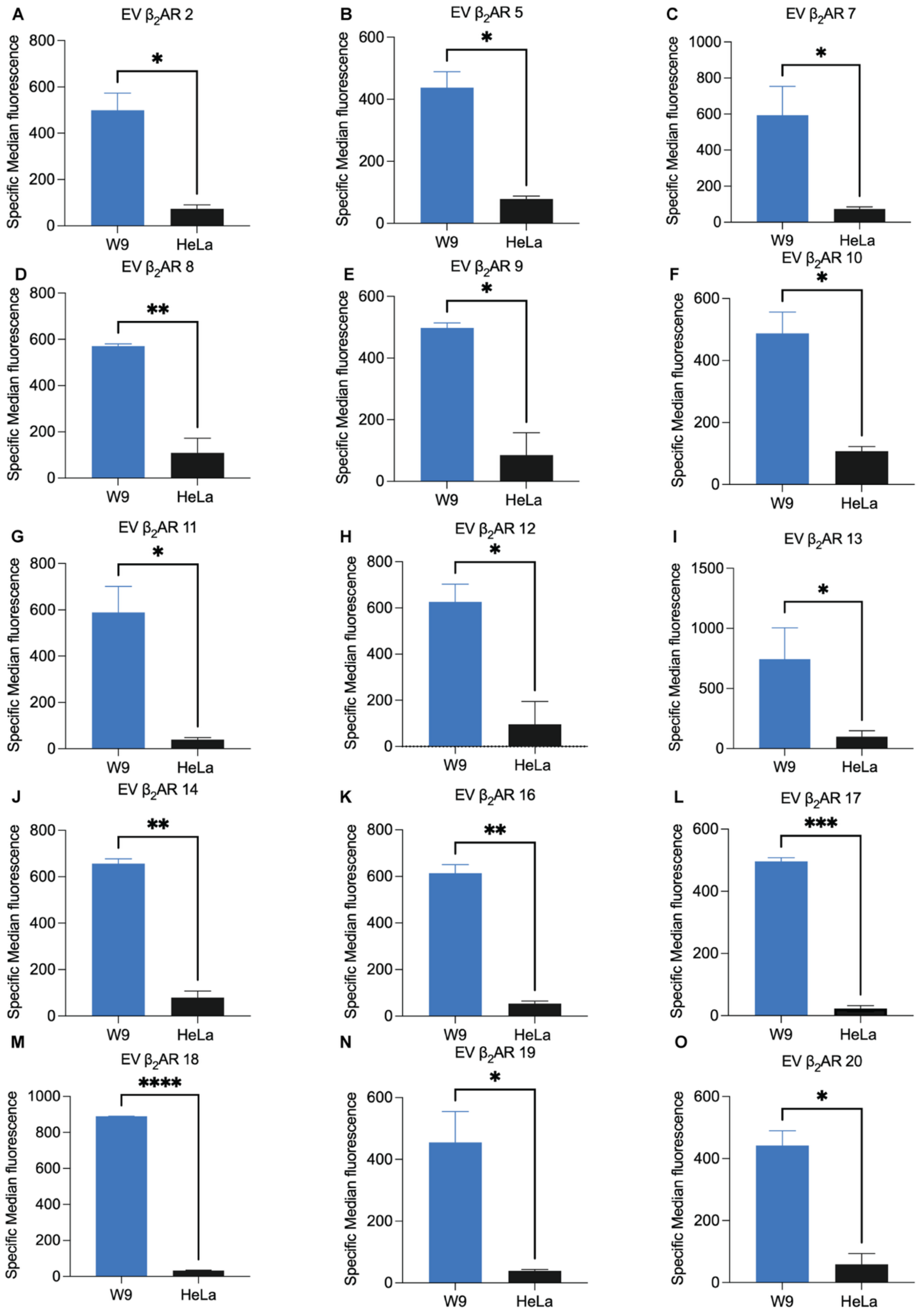

A
EV β2AR 2
B
EV β2AR 5
C
EV β2AR 7
D
EV β2AR 8
E
EV β2AR 9
F
EV β2AR 10
G
EV β2AR 11
H
EV β2AR 12
I
EV β2AR 13
J
EV β2AR 14
K
EV β2AR 16
L
EV β2AR 17
M
EV β2AR 18
N
EV β2AR 19
O
EV β2AR 20
Specific Median fluorescence
W9
HeLa

**Figure S13: The binding of hit aptamer candidates to W9 cells.** A-O) Binding of the selected hit aptamer candidates to W9 cells was evaluated by incubating $1.5 \times 10^5$ W9 or HeLa cells with 500 nM of the fluorescently labeled hit aptamer candidate or a random sequence at 4 °C for 45 minutes. Subsequently, binding interactions were quantified through flow cytometry, and the median fluorescence intensity was used to calculate the specific median fluorescence according to the following equation: specific median fluorescence = [median fluorescence of aptamer candidate − median fluorescence of random]. Fluorescence intensity measurements indicate strong binding affinity to W9 cells overexpressing the target with minimal signal detected in HeLa cells. The specific median fluorescence of each candidate was graphically presented using GraphPad Prism software and further analyzed via a non-parametric *t*-test. Statistical significance was established at a *p*-value of 0.05.

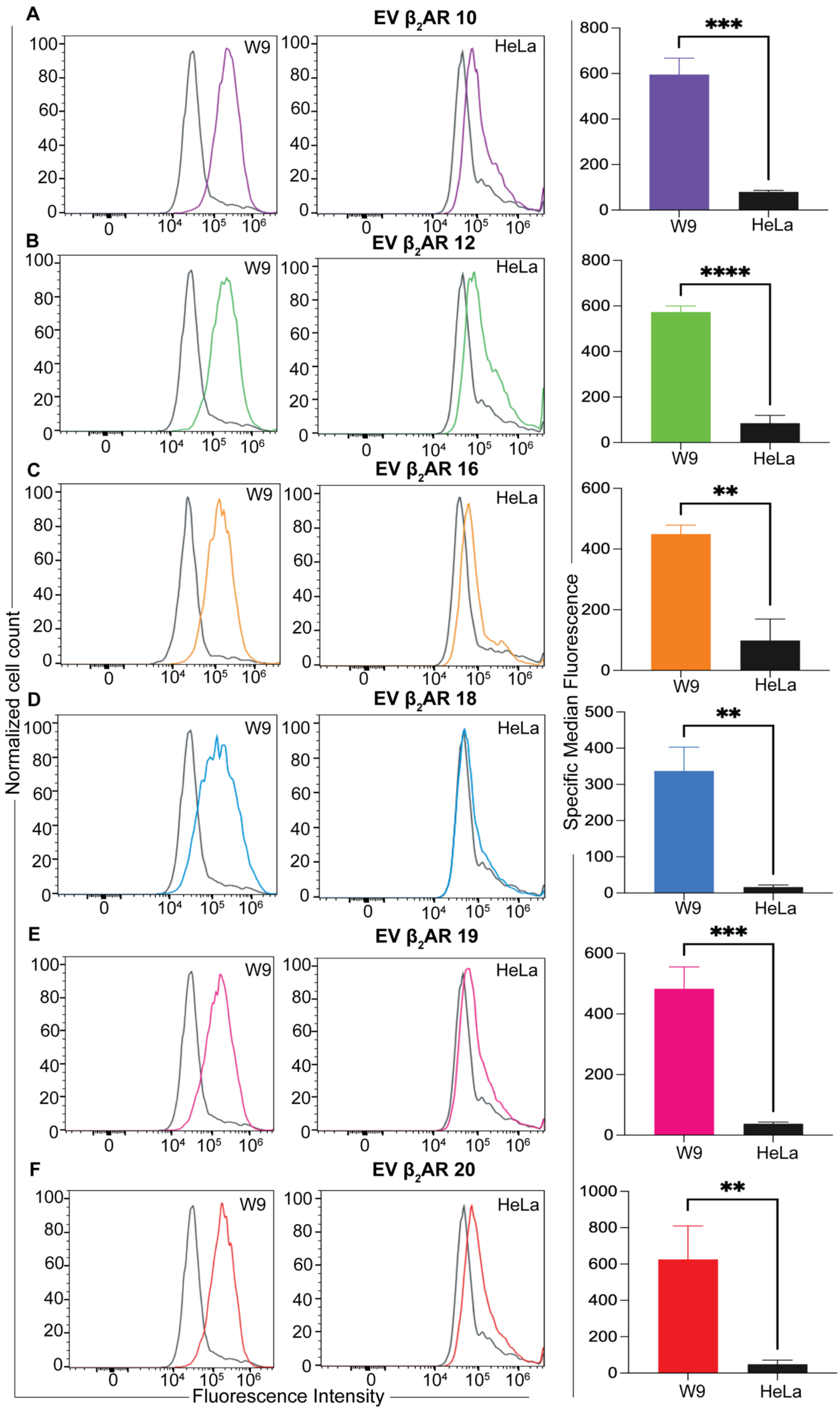
A
EV β2AR 10
W9
HeLa
B
EV β2AR 12
C
EV β2AR 16
D
EV β2AR 18
E
EV β2AR 19
F
EV β2AR 20
Normalized cell count
Fluorescence Intensity
Specific Median Fluorescence
***
****
**
**
***
**

**Figure S14: Specificity of hit aptamer candidates for binding to W9 cells at 37 ºC.** A-F) The binding of hit aptamer candidates to W9 cells was examined by incubating 7.5 × $10^4$ W9 or HeLa cells with 250 nM of the fluorescently labeled hit aptamer candidate, or a random sequence, at 37 °C for 45 minutes. Subsequently, binding was quantified through flow cytometry, and the median fluorescence was employed to calculate the specific median fluorescence according to the following equation: specific median fluorescence = [(median fluorescence of aptamer candidate − median fluorescence of random)/ median fluorescence of random] x 100. The median fluorescence of each candidate was plotted using GraphPad Prism software and further analyzed using a non-parametric *t*-test; the data represent the mean ± SD of three independent experiments. Fluorescence intensity measurements demonstrate strong binding to W9-overexpressing cells with minimal signal detected in HeLa cells.

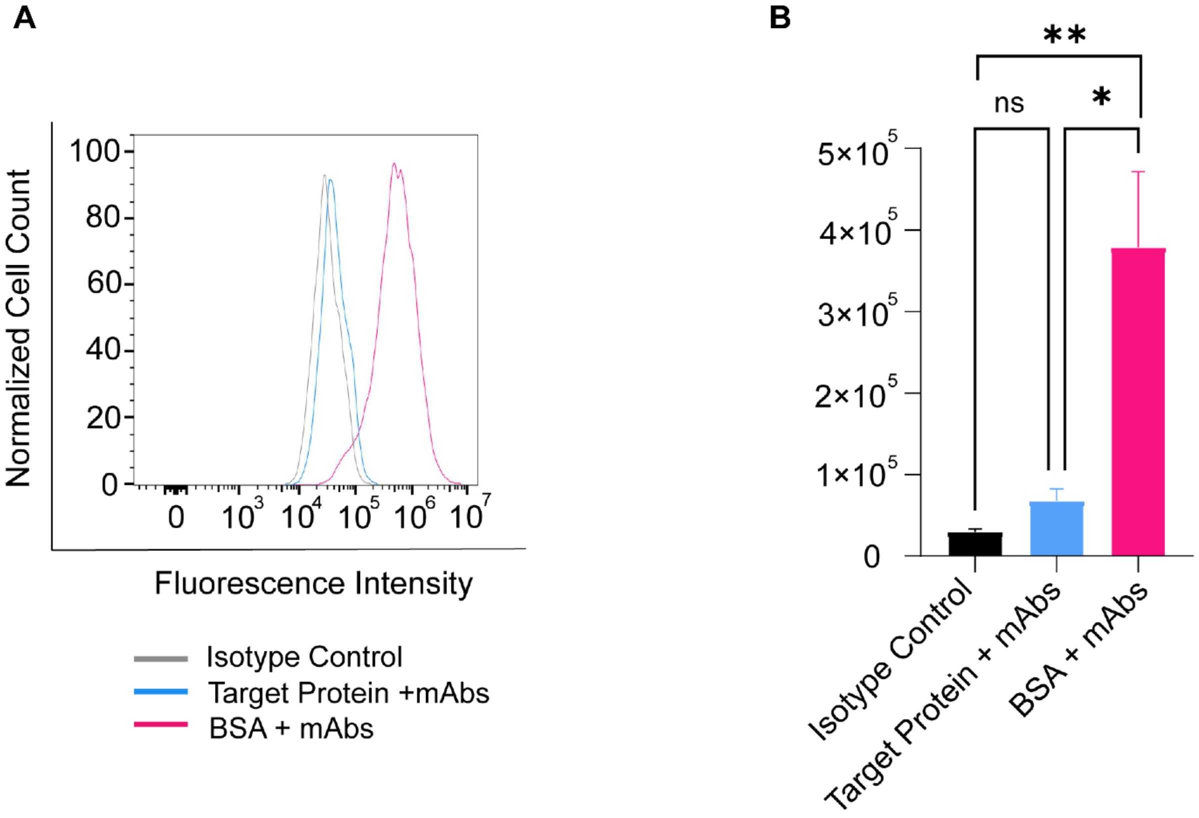


**Figure S17: Competition binding assay of FLAG mAbs to $\beta_2$AR-expressing W9 cells.** A) Representative flow cytometry histograms showing FLAG mAbs binding to W9 cells under different pretreatment conditions. Cells pretreated with purified $\beta_2$AR and incubated with isotype control (gray) displayed minimal fluorescence, serving as negative control. Pretreatment with $\beta_2$AR followed by incubation with FLAG mAbs resulted in a pronounced leftward shift in fluorescence intensity (blue), indicating reduced antibody binding owing to receptor competition. In contrast, cells pretreated with BSA and incubated with FLAG mAbs (pink) exhibited higher fluorescence intensity, reflecting robust antibody binding in the absence of competition. (b) Bar graph illustrating the average median fluorescence intensity (MFI) from three independent experiments (n=3), as represented by mean ±

SEM. Pretreatment with purified β2AR significantly reduced the binding of FLAG mAbs (blue) compared to BSA-treated cells (pink) ($p$< 0.05, one-way ANOVA).

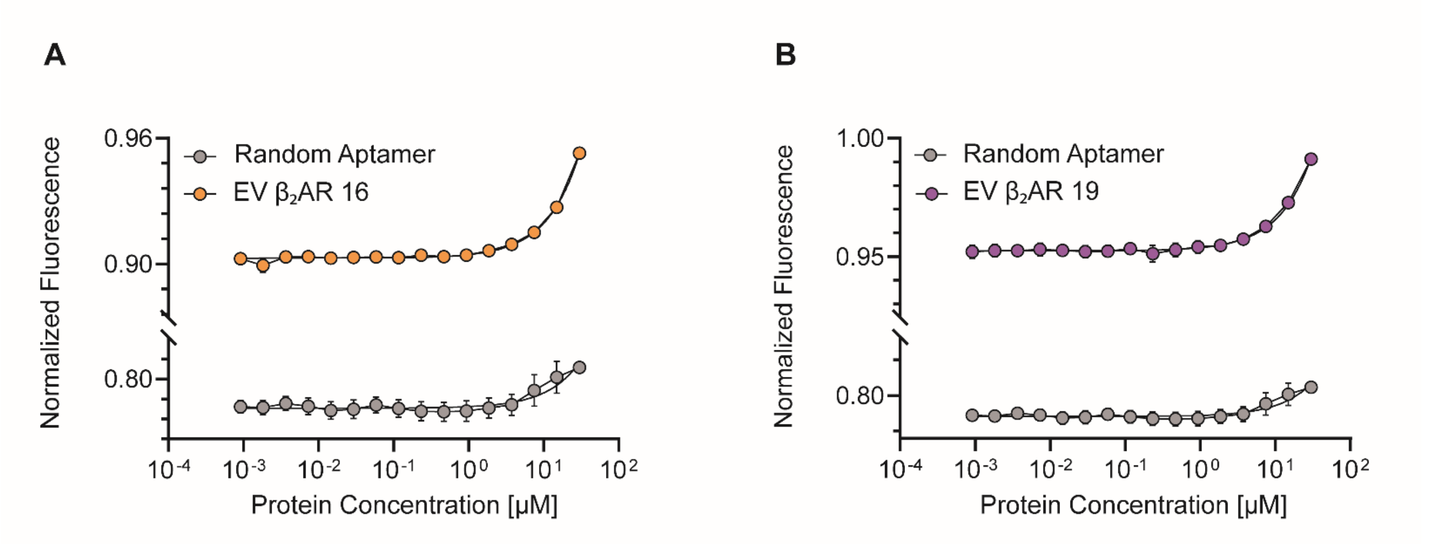


**Figure S18: Representative MST normalized fluorescence of Cy5-labeled aptamers in the presence of unlabeled β2AR and BSA.** A fixed aptamer concentration of 1μM was titrated with increasing concentration of the purified protein (9.1 nM to 30 μM) and random as a negative control. The X-axis shows increasing protein concentration, and the Y-axis shows normalized fluorescence measure on the Monolith X instrument (NanoTemper). Data represented are mean ± SEM from three independent experiments (n=3). Statistical significance was determined using XY linear regression analysis ($p$-value < 0.0001).

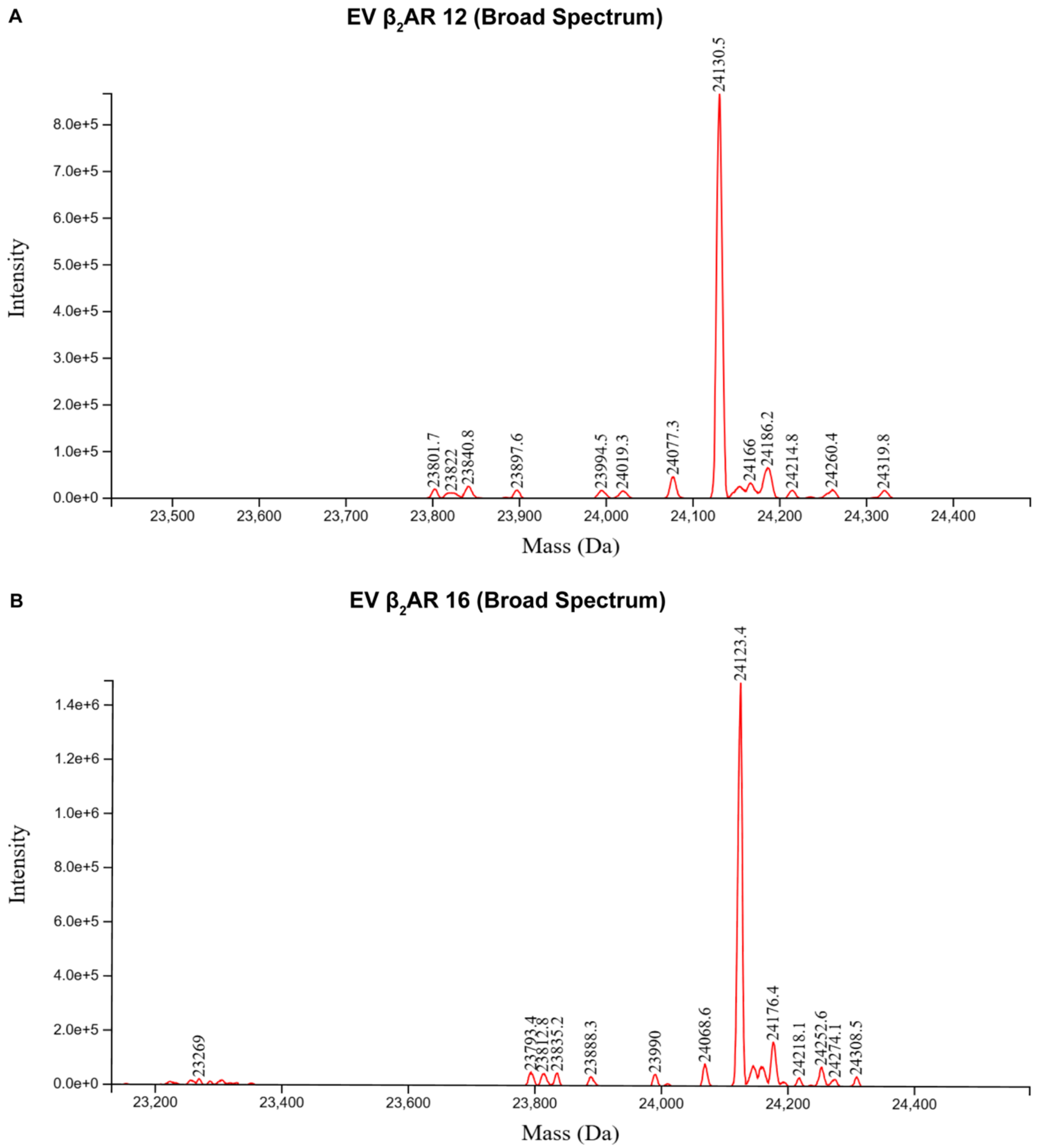


**Figure S19: LC-MS characterization of synthesized $\beta_2$AR DNA aptamers.** Deconvoluted electrospray ionization liquid chromatography–mass spectrometry (LC–ESI–MS) spectra were used to evaluate the molecular masses of two EV $\beta_2$AR DNA aptamer sequences with an expected length of 77 nucleotides. The calculated molecular masses for the full-length aptamers are 23,562.2 Da for EV $\beta_2$AR 12 and 23,555.2 Da for EV $\beta_2$AR 16. In both cases, the dominant observed peaks exhibited a mass increase of approximately +568 Da relative to the calculated masses (+568.3 Da for EV $\beta_2$AR 12 and +568.2 Da for EV $\beta_2$AR 16), consistent with the presence of n+2 extended species (~79 nt).

These results indicate that the major products correspond to aptamer sequences containing two additional nucleotides compared with the expected 77-nt length.

**Table S5: Calculation of the molecular masses for the full-length $\beta_2$AR aptamers.**

| Molecules Names | IDT Mass Calculated [Da] | Mass Found [Da] | Major Peak [Da] |
|---|---|---|---|
| EV $\beta_2$AR 12 | 23562.2 | 24130.5 | 24130.5 |
| EV $\beta_2$AR 16 | 23555.2 | 24123.4 | 24123.4 |

1. EV $\beta_2$AR 12: extra 568.3 Da meaning n+2 (~79nt-mer)
2. EV $\beta_2$AR 16: extra 568.2 Da meaning n+2 (~79nt-mer)
